# Feasibility of 3T layer-dependent fMRI with GE-BOLD using NORDIC and phase regression

**DOI:** 10.1101/2022.06.02.494602

**Authors:** Lasse Knudsen, Christopher J. Bailey, Jakob U. Blicher, Yan Yang, Peng Zhang, Torben E. Lund

## Abstract

**Introduction:** Functional MRI with spatial resolution in the submillimeter domain enables measurements of activation across cortical layers in humans. This is valuable as different types of cortical computations, e.g., feedforward versus feedback related activity, take place in different cortical layers. Layer-dependent fMRI (L-fMRI) studies have almost exclusively employed 7T scanners to overcome the reduced signal stability associated with small voxels. However, such systems are relatively rare and only a subset of those are clinically approved. In the present study, we examined the feasibility of L-fMRI at 3T using NORDIC denoising.

**Methods:** 5 healthy subjects were scanned on a Siemens MAGNETOM Prisma 3T scanner. To assess across-session reliability, each subject was scanned in 3-8 sessions on 3-4 consecutive days. A 3D gradient echo EPI (GE-EPI) sequence was used for BOLD acquisitions (voxel size 0.82 mm isotopic, TR = 2.2 s) using a block designed finger tapping paradigm. NORDIC denoising was applied to the magnitude and phase time series to overcome limitations in tSNR and the denoised phase time series were subsequently used to correct for large vein contamination through phase regression.

**Results and conclusion:** NORDIC denoising resulted in temporal signal-to-noise ratio (tSNR) values comparable to or higher than commonly observed at 7T. Layer-dependent activation profiles could thus be extracted robustly, within and across sessions, from regions of interest located in the hand knob of the primary motor cortex (M1). Phase regression led to substantially reduced superficial bias in obtained layer profiles, although residual macrovascular contribution remained. We believe the present results support the feasibility of L-fMRI at 3T, which might help make L-fMRI available to a much wider community.

## 1. Introduction

Functional MRI (fMRI) with spatial resolutions at the submillimeter scale is a rapidly growing field, motivated, in part, by the ability to resolve cortical layers noninvasively in human subjects. Activation patterns across distinct laminae constitute a fingerprint of feedforward and feedback related information (Felleman & Van Essen, 1991), making the method, which we will refer to as L-fMRI, a valuable asset for studying hierarchical information flow in the brain. Numerous studies already demonstrated its worth in areas related to, e.g., visual perception and attention (Aitken et al., 2020; Kok et al., 2016; Liu et al., 2020; Muckli et al., 2015), motor control (Huber et al., 2017; Persichetti et al., 2020), somatosensation (Yu et al., 2019, 2022), auditory perception (De Martino et al., 2015; Moerel et al., 2019), and language (Sharoh et al., 2019). L-fMRI further holds promise as a tool to expand our understanding of the diseased brain, as proposed specifically for neurodegenerative diseases, in a recent review (McColgan et al., 2020).

Resolving functional responses of such delicate structures is no easy task and comes with several challenges (Bandettini et al., 2021; Polimeni et al., 2018), a major one being reduced tSNR resulting from the small voxel sizes, which is magnified by the fact that regular spatial smoothing is prohibited as it would wash out the layer-specific information. Another concern is the well-established degradation of spatial specificity caused by blood-signal being dispersed over relatively large distances (Turner, 2002) towards and along the cortical surface (Menon, 2012; Uludağ et al., 2009). Ascending veins and pial veins constitute the cortical infrastructure underlying this drainage (Duvernoy et al., 1981) and the problem applies in particular to BOLD fMRI using GE-EPI readouts (GE-BOLD) due to its T2* based contrast (Yacoub et al., 2003). Sequences relying on T2 based contrast such as spin echo EPI (SE-EPI) (Han et al., 2021; Koopmans & Yacoub, 2019) and b-SSFP (Báez-Yánez et al., 2017; Liu et al., 2020) or cerebral blood volume (CBV) based contrast such as VASO (Huber et al., 2015; Lu et al., 2003) are less sensitive to large veins and therefore have better spatial specificity. However, SE-EPI for instance is rarely free of T2* effects due to long readout times (Goense & Logothetis, 2006) and trade-offs with these alternatives to GE-BOLD generally include lower contrast-to-noise ratio (CNR), lower sampling efficiency and potentially less straightforward implementation. The “best” sequence thus doesn’t exist and which one to choose depends on the specific research goal. GE-BOLD has nevertheless been the most frequently adopted sequence for L-fMRI hitherto and will be the sequence of focus in the present study.

The vast majority of L-fMRI studies employed ultrahigh field (≥7T) scanners which is motivated by an accompanied increase in tSNR (Triantafyllou et al., 2005) plus an enhanced BOLD effect (Ugurbil, 2014). Moreover, signal contributions from intravascular compartments are reduced as a result of the very short T2-value of blood at ultrahigh field (Jochimsen et al., 2004; Uludağ et al., 2009). Ultrahigh field hence improves both the sensitivity and specificity problems (Chaimow et al., 2018; Dumoulin et al., 2018; Uğurbil, 2021). The downside of a dependence on ultrahigh field is a dramatic decrease in the availability of L-fMRI, as these systems are still relatively rare, and clinically approved ones, even more so. Successful GE-BOLD 3T implementations of L-fMRI have been reported (Koopmans et al., 2010; Markuerkiaga et al., 2020; Puckett et al., 2016; Ress et al., 2007; Scheeringa et al., 2016), demonstrating that useful layer-dependent activation measures can be obtained outside of ultrahigh field applications. However, the sequences and analysis strategies adopted in these studies to overcome the reduced sensitivity at lower field strength may not be generally applicable and widespread use of L-fMRI at 3T has thus yet to emerge. To this end, a recently published denoising method, named NORDIC (Moeller et al., 2021; Vizioli et al., 2021), has proven effective at increasing the signal stability of magnitude and phase time series data. It is based on principal component analysis (PCA) applied to the full complex valued MRI dataset data with the aim of removing principal components that cannot be distinguished from zero-mean gaussian noise. Thermal noise fits this description and is the dominant source of noise for submillimeter voxels at 3T (Triantafyllou et al., 2005). Initial reports on the effect of NORDIC indicate substantial tSNR increments without any significant spatial blurring being introduced (Dowdle et al., 2021; Vizioli et al., 2021). It may thus help moving the field one step closer towards overcoming the problem of low tSNR which currently limits routine use of L-fMRI at 3T.

Accordingly, the purpose of this study is to examine the feasibility of 3T L-fMRI with GE-BOLD using NORDIC denoising. Furthermore, as pointed out in the above, the choice of GE-BOLD entails large vein correction for which a number of post processing approaches have been proposed (discussed in Huang et al., 2021; Kay et al., 2019; Stanley et al., 2020). Here we examine the approach referred to as phase regression (see Menon, 2002 and Methods section 2.4.2) for two main reasons: 1) it has proved effective for selectively suppressing the response in voxels contaminated by large veins (pial veins and largest ascending veins) while maintaining high sensitivity (Stanley et al., 2020), 2) It appears to be available for a large range of L-fMRI applications as it doesn’t rely on any additional scans or equipment, and it doesn’t, in principle, rely on paradigm design (although it has some CNR dependency). We scanned subjects during a finger tapping task and extracted layer profiles from regions of interest (ROIs) in the hand knob of the primary motor cortex (M1) which enabled comparison with results from the 7T L-fMRI literature. Finally, we examined the reliability of this 3T setup both within and across sessions by scanning each subject on multiple days. The results could potentially help to increase the utility of L-fMRI at 3T, which would be valuable for making L-fMRI available to a much wider community.

## 2. Methods

### 2.1 Subjects

5 healthy right-handed subjects (1 female) with an age of 25-29 years were included in the study, approved by the regional ethics committee in Region Midt (Study ID 1-10-72-216-21). All subjects were carefully informed about the procedures and provided written consent.

### 2.2 Imaging protocol

Imaging was performed using a Siemens MAGNETOM Prisma 3 T scanner equipped with a standard 32Ch-receive head coil. Anatomical reference images were collected with a MP2RAGE sequence (Marques et al., 2010) and parameters: voxel size = 0.9 mm isotropic, matrix size = 192 × 240 × 256, iPAT = 2, Partial Fourier = 6/8, TE = 2.87 ms, TR = 5000 ms, TI1 = 700 ms, FA1 = 4°, TI2 = 2500 ms, FA2 = 5°, echo spacing = 7.18 ms. Functional images were obtained with a 3D GE-EPI sequence (Stirnberg & Stöcker, 2021) and parameters: voxel size = 0.82 mm isotropic, TR = 2200 ms, TE = 27 ms, iPAT = 3, Partial Fourier = 6/8 (zero-filling reconstruction), FA = 45 degrees, and matrix size = 176 × 176 × 26 where the 26 axial slices were aligned perpendicularly with respect to the surface of the hand knob in M1 (Figure 1). Magnitude and phase reconstruction was done using adaptive coil combine (Walsh et al., 2000). Additionally, we used a SS-SI-VASO sequence (Huber et al., 2014) (voxel size = 0.82 mm isotropic, TR = 4739 ms, TE = 27 ms, iPAT = 3, Partial Fourier = 6/8, FA = 18 degrees, and matrix size = 176 × 176 × 26, volumes = 100) with the same readout strategy and resolution as the functional sequence to obtain T1-weighted images with good anatomical contrast, and distortions similar to the functional images. Matched distortions enabled high quality registration to the functional images without the need for distortion correction, and the T1-weighted contrast further aided nonlinear registration of the MP2RAGE image to EPI volumes.

**Figure 1.**
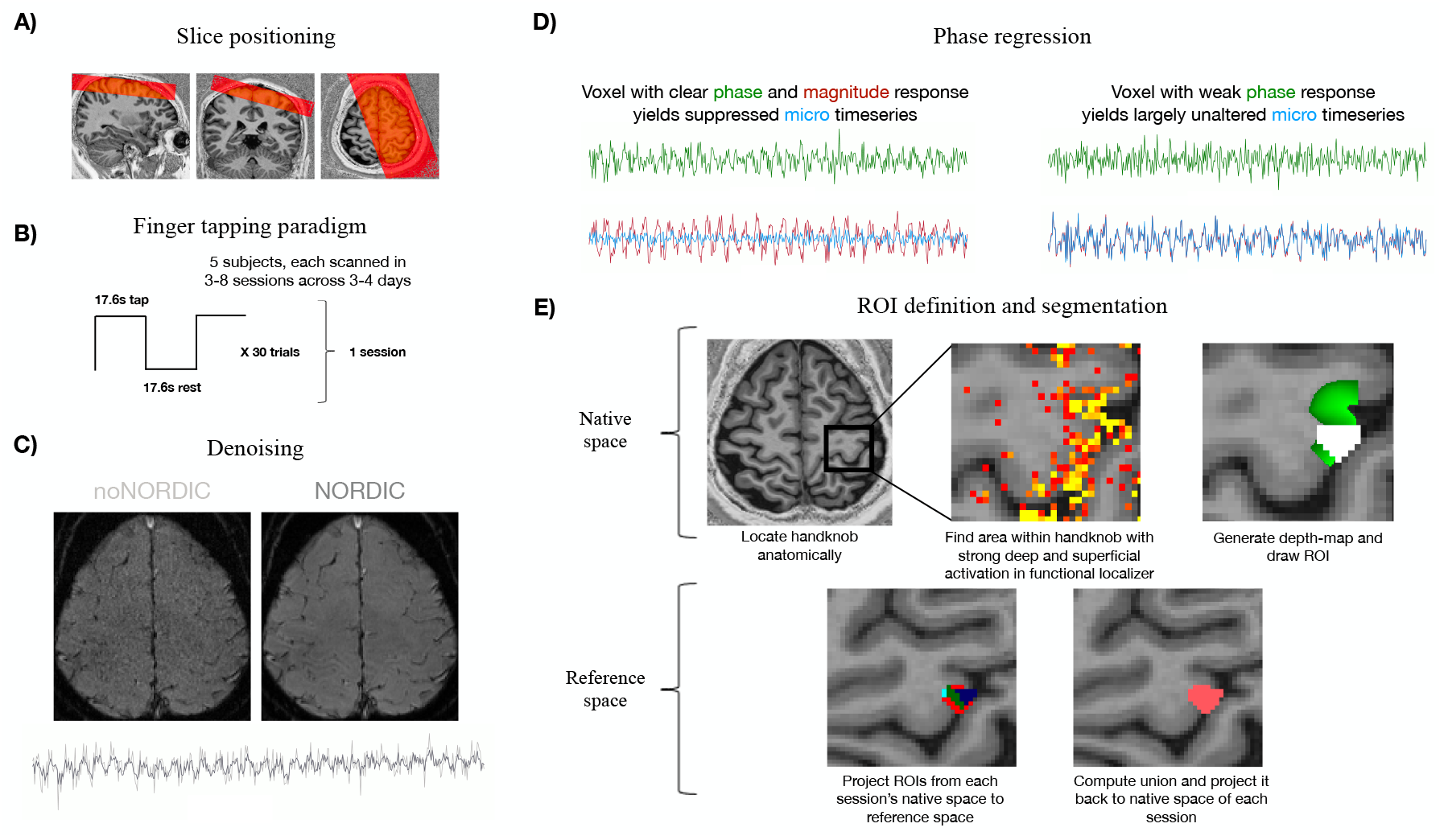
Overview of acquisition and analysis pipeline **A)** The imaging slab was positioned approximately perpendicular to the cortical surface of the hand knob in M1 (left hemisphere) based on the initially acquired anatomical image (MP2RAGE). **B)** The functional paradigm of each session consisted of ∼18 minutes (including functional localizer) block designed finger tapping (17.6 s tapping alternating with 17.6 s rest, repeated in 30 trials). **C)** NORDIC denoising was applied to the functional timeseries data after DICOM-to-NIFTI conversion. **D)** Motion corrected NORDIC and noNORDIC timeseries were then submitted to phase regression to correct for large vein effects. An example phase timeseries from a voxel which is located towards CSF, shown in green to the left, has a clear modulation with respect to the task and thus presumably contains a large vein. The linear fit between the phase timeseries and magnitude timeseries is subtracted from the original magnitude signal (red) leading to suppression of the paradigm modulation in the resulting “micro” timeseries (blue). The same is shown to the right for a voxel in gray matter where a clear task modulation can be observed in the magnitude timeseries only, resulting in a largely unchanged micro timeseries. **E)** ROIs of each subject were defined in native space by first anatomically locating the hand knob in each session (Yousry et al., 1997). They were then manually drawn in the part of the knob where all sessions had strong activation in the functional localizer. Next, ROIs from each session were aligned to reference space (first session of each subject) and a common union ROI was computed and projected back to native space of each session.

### 2.3 Experimental protocol

Each subject was scanned in 3-8 sessions on 3-4 consecutive days resulting in 23 total sessions, 3 of which were excluded due to excessive motion or artifacts. The procedure was identical across all sessions to enable assessment of across-session consistency of laminar responses. The paradigm consisted of a single functional run with 35.2 s trials (17.6 s tapping and 17.6 s rest) repeated 30 times resulting in a total duration of ∼18 minutes and acquisition of 480 functional volumes (exclusive 4 dummy volumes). Tapping blocks included alternating finger tapping movements between the right index and middle fingers. The frequency of tapping for each finger was ∼2.5 Hz and was guided visually by a blinking marker projected onto a semi-transparent screen placed inside the MRI scanner, visible to the subject via a mirror mounted on the head coil. Before entering the scanner, subjects were instructed to lie as still as possible, while keeping their gaze on the blinking marker in both tapping and resting blocks. Head motion was further minimized by tape across the forehead for haptic feedback and by placing inflatable padding around the head which could be manually adjusted by the subject to optimize comfort. Body motion was reduced by wrapping the subject in a heavy blanket. The duration of each scanning session was kept below 1 hour.

### 2.4 Data analysis

#### 2.4.1 Preprocessing

After DICOM-to-NIFTI conversion, functional and VASO time series were denoised with NORDIC (Vizioli et al., 2021, https://github.com/SteenMoeller/NORDIC_Raw, downloaded 27042022). In NORDIC, singular value decomposition (SVD) is performed on patches of functional time series data (represented as Casorati matrices), where all components with singular values below a certain threshold are removed. This threshold is determined as the average of the largest singular value across Monte Carlo-simulated thermal noise matrices. The variance associated with these random matrices is estimated directly from the measured data, either by an appended noise-only acquisition or from an area outside the brain without signal contributions (see Vizioli et al., 2021 for details). Here, the latter approach was employed. In addition, we employed more aggressive denoising by setting the option ARG.factor_error (FE) equal to 1.15 (default is 1). The more aggressive denoising comes with a risk of altering the temporal correlations in the data, but as shown in Figure S1 and Figure S2, additionally removed components appeared to be dominated by thermal noise.

The denoised functional magnitude time series were then motion corrected in SPM12 (Functional Imaging Laboratory, University College London, UK). To optimize the alignment around the hand knob of M1, a spatial weighting mask was applied with largest weights on the hand knob and progressively smaller weights towards the periphery of the brain. The phase images were converted into real and imaginary parts prior to reslicing using realignment parameters estimated from the magnitude images. These resliced (complex) images were converted back to phase values and unwrapped in the temporal direction. For the purpose of comparing scenarios with and without application of NORDIC, the estimated motion parameters were also applied to the same magnitude and phase time series without denoising (noNORDIC).

SS-SI-VASO provides 2 images per TR, one where the blood has been nulled by an inversion pulse, and a second where the blood has not been nulled (Huber et al., 2014). These images were first motion corrected separately due to their distinct contrasts, and then registered to the first image of the functional time series in a single interpolation step. The inverse signal variability of the combined nulled and not-nulled time series was then used to compute a T1-weighted image with high anatomical contrast (Beckett et al., 2020; Huber et al., 2017). Due to this VASO EPI image theoretically being in the same distorted space as the functional BOLD EPI images while having similar contrast as MP2RAGE, it was used as a “high-quality reference image” for non-linear registration of MP2RAGE to EPI-space. This was done by first estimating initial parameters for alignment of MP2RAGE to the T1-weighted reference in ITK-SNAP (Yushkevich et al., 2006), followed by rigid, affine and non-linear registration steps (*SyN* algorithm) in ANTS (Avants et al., 2011). We carefully inspected the outputs to ensure all registrations were of sufficient accuracy for laminar analysis.

#### 2.4.2 Implementation of phase regression

Next, phase regression was applied according to the scheme described in previous reports (Curtis et al., 2014; Menon, 2002; Stanley et al., 2020; Vicente et al., 2021) to correct for bias associated with large veins. The main principle is to remove the linear fit between magnitude and phase time series from the magnitude time series. This is based on the physical property that macrovascular directional flow changes result in a change in the phase signal, whereas microvascular flow has no common direction resulting in no net phase change (for details, see Menon, 2002). We will refer to the resulting time series as the micro time series. Before phase regression, magnitude and phase time series were filtered on a voxel wise basis by first fitting a general linear model (GLM) with a 16’th order FIR-set modeling the paradigm, and 60 nuisance regressors (24 motion (Friston et al., 1996), 20 RETROICOR (Glover et al., 2000) and 16 high pass) and then subtracting the nuisance fit from the original time series. Prior to nuisance regression the design matrix containing the nuisance regressors was orthogonalized with respect to the paradigm. In the event of stimulus locked motion, this orthogonalization leaves paradigm effects in the phase regressor unchanged even after nuisance filtering. Voxel wise linear fit parameters between filtered magnitude and phase time series were estimated with a deming regression model (Hall, 2022), which is necessary as it accounts for noise on both the x- and y-variables, whereas standard Ordinary Least Squares (OLS) regression assumes noise free regressors. The fit needs to be conditioned by the relative noise levels between magnitude and phase data, which determines the angle of the residuals to the line of the best fit. To accommodate this, the voxel-wise residual variance in the filtered magnitude and phase time series was estimated by further subtracting the fit of the 16’th order FIR-set to also remove the variance explained by the paradigm. The relative noise level was then given as the ratio between the standard deviation of the magnitude residuals and the standard deviation of the phase residuals.

#### 2.4.3 Activation maps

GLMs were then computed using AFNI (Cox, 1996) where the design matrix consisted of a single paradigm regressor convolved with a canonical hemodynamic response function, which was done separately for each of the following time series; filtered magnitude time series with and without NORDIC, and micro time series with and without NORDIC, resulting in 4 statistical maps per session (called NORDICmagn, noNORDICmagn, NORDICmicro and noNORDICmicro, respectively). Time series and regressors were scaled to yield beta maps reflecting percent signal change with respect to baseline. A separate parameter estimate (beta) was estimated for each of the 30 tapping trials and odd trials were used for functional localization of ROIs only (unless stated otherwise), whereas even trials were used in all further analyses. In order to minimize the impact of large veins during localization, the beta values for odd trials were extracted from time series denoised with FE = 1.4 whereas beta values for even trials were extracted from the same time series but denoised with FE = 1.15.

#### 2.4.4 ROI definition and segmentation

ROIs were manually defined in the hand knob area of Brodmann area 4a (BA4a) similarly to previous layer-fMRI reports focusing on M1 and finger tapping (Beckett et al., 2020; Chai et al., 2020; Han et al., 2021; Huber et al., 2017; Persichetti et al., 2020; Shao et al., 2021). First, the hand knob was visually identified from its characteristic omega-shape (Yousry et al., 1997) and voxel selection was then limited to the upper and lateral part of the knob corresponding to BA4a (Huber et al., 2017). For identification of the index-middle finger representation, we found the area which had strong localizer activation in both deep and superficial layers across all sessions for that subject. Gray matter (GM) boundaries were manually drawn on an upsampled grid (0.2 mm in plane resolution) around this area guided by an automated segmentation obtained with CAT12 (http://www.neuro.uni-jena.de/cat/). A depth map comprising the relative cortical depth (values between 0 and 1) of voxels between these boundaries was then computed with LAYNII (Huber et al., 2021) using the equivolume metric (Waehnert et al., 2014). The point at which voxels were estimated to be 50/50 white matter (WM) and GM was located at a relative depth between 0.1 and 0.2, and the equivalent point for GM and cerebrospinal fluid (CSF) was between 0.8 and 0.9. After segmentation, the set of voxels constituting the ROI was selected to maximize the signal in depths 0.3-0.4 (deep layers). 0.3 was chosen as lower bound to minimize partial voluming with WM and 0.4 was chosen as upper bound due to expected signal reductions in middle layers (Chai et al., 2020; Huber et al., 2017; Shao et al., 2021). This selection was restricted by the following criteria: 1) ROIs were defined as columns (no holes) going roughly perpendicular onto the surface to make it somewhat anatomical meaningful; 2) ROIs had to consist of minimum 50 voxels to increase the ability to compare the same cortical portion across sessions despite e.g. imaging slab positioning not being fully identical, to minimize influence of inaccuracies in e.g., segmentation or across session alignment, and to increase representability of measured M1 activation; 3) ROIs should not extend into more than four slices; 4) the peak signal in depths 0.3-0.4 should not exceed that of depths 0.6-0.8 in order to identify the same functional representation as previous M1 layer-fMRI studies, which generally found stronger activity in superficial compared to deep layers (Beckett et al., 2020; Chai et al., 2020; Han et al., 2021; Huber et al., 2017; Shao et al., 2021); 5) Empty axial slices were not allowed in the final union ROI (see below), that is, it had to be consecutive. These initial ROIs defined in native space of each individual session were then transformed into the space of each subject’s first session where the union of all ROIs was computed and transformed back to native space (Figure 1). Union ROIs were manually corrected in native space in cases with obvious anatomical mistakes, such as holes in the middle of a column. The initial native ROIs were solely used to compute the union-ROI from which data for all further analysis was extracted. This approach was adopted to ensure the same area of the knob was compared across all sessions for each subject.

#### 2.4.5 Generation of layer profiles

Layer profiles were generated by plotting the beta of each voxel within the ROI as a function of its cortical depth. Notice that beta-maps were kept in their original resolution and the depth of a given voxel, that is, the correspondence between its center and the underlying depthmap in upsampled resolution, was inferred by Nearest Neighbor interpolation using the *spm_sample_vol*-function in SPM12. When plotting beta-estimates from single voxels, the resulting cloud of data points becomes quite noisy, so MATLAB’s (Mathworks Inc.) *movmean*-function (window size = 0.2) was used to get the final layer profiles for each of the 15 test-betas which were averaged to obtain a single profile per session per data version. In order to calculate mean profiles across sessions and subjects, without bias from specific depths being sampled more in some sessions or subjects, profiles were interpolated at 18 equally spaced depths between 0.075 and 0.925. This approach was chosen, over interpolation of the activation maps, in order to more directly interpolate between voxels sampled at similar depths.

#### 2.4.6 Sensitivity and specificity analysis

Sensitivity analysis was conducted by submitting the 15 betas to a second level analysis where voxel wise t-values were calculated, then averaged within the ROI for each session, and finally averaged across sessions to get a single t-value per subject per data version. This approach is in contrast to the typical calculation of t-values from a first level fMRI analysis where residuals from the GLM is used directly. This approach was chosen to acknowledge that the degrees of freedom potentially may be affected by NORDIC denoising. By generating t-values from single trial beta-estimates, which are assumed to be independent, this potential concern is largely removed.

A quantitative measure of spatial specificity was obtained by fitting a line to the across-session mean layer profiles of each subject. The slopes of these lines represent the degree of bias towards the pial surface, i.e. inverse specificity (Beckett et al., 2020). Two-way repeated measures ANOVAs with factors *NORDICversion* (NORDIC/noNORDIC) and *veinCorrection* (micro/magnitude) were employed to evaluate statistical significance of differences in sensitivity (t-values) and specificity (slopes) across the 4 combinations of NORDIC/noNORDIC and magnitude/micro data versions.

Data is presented as mean ± standard deviation unless otherwise stated. The significance level was set to α= 0.05.

### 2.5 Data and code availability

The data are available upon request and signed data sharing and data processing agreements as part of Aarhus University data sharing regulations. Analysis code will be made publicly available on figshare: https://doi.org/10.6084/m9.figshare.19925483.v1

## 3. Results

NORDICmicro layer profiles for each of the 15 test trials are shown for each session and subject in Figure 2. Laminar patterns of activation appear highly consistent across trials, with trial profiles closely following the mean profiles of individual sessions. Similarly to what was done in Huber, Tse, et al., (2018), we obtained a quantitative measure of this reproducibility by calculating the coefficient of variation (CV), i.e., standard deviation across trials divided by the mean, which was done for each layer and session separately and then averaged. The average trial-CV across subjects was 20.84 % ±3.70 %. Consistency across trials was clearly facilitated by NORDIC, which is evident from Figure 3A where NORDICmagn and noNORDICmagn profiles are compared in a representative subject. Across-trial mean profiles are practically identical whereas the variability around the mean is notably increased when denoising is omitted. Within session stability was further quantified by computing an average tSNR-value of all ROI-voxels for each session which was averaged to one tSNR-value per subject. This was done using detrended but otherwise unfiltered motion corrected magnitude time series with and without NORDIC. The mean tSNR was ∼3 times larger with NORDIC (39.47 ±5.07), compared to without NORDIC (12.28 ±1.37) (paired t-test, degrees of freedom = 4, p < 0.001). The effect of such increased temporal stability on activation maps is illustrated in an example subject in Figure 3B. In the high tSNR case, the hand knob is clearly activated and accompanied by much less false positive activation outside of M1.

**Figure 2.**
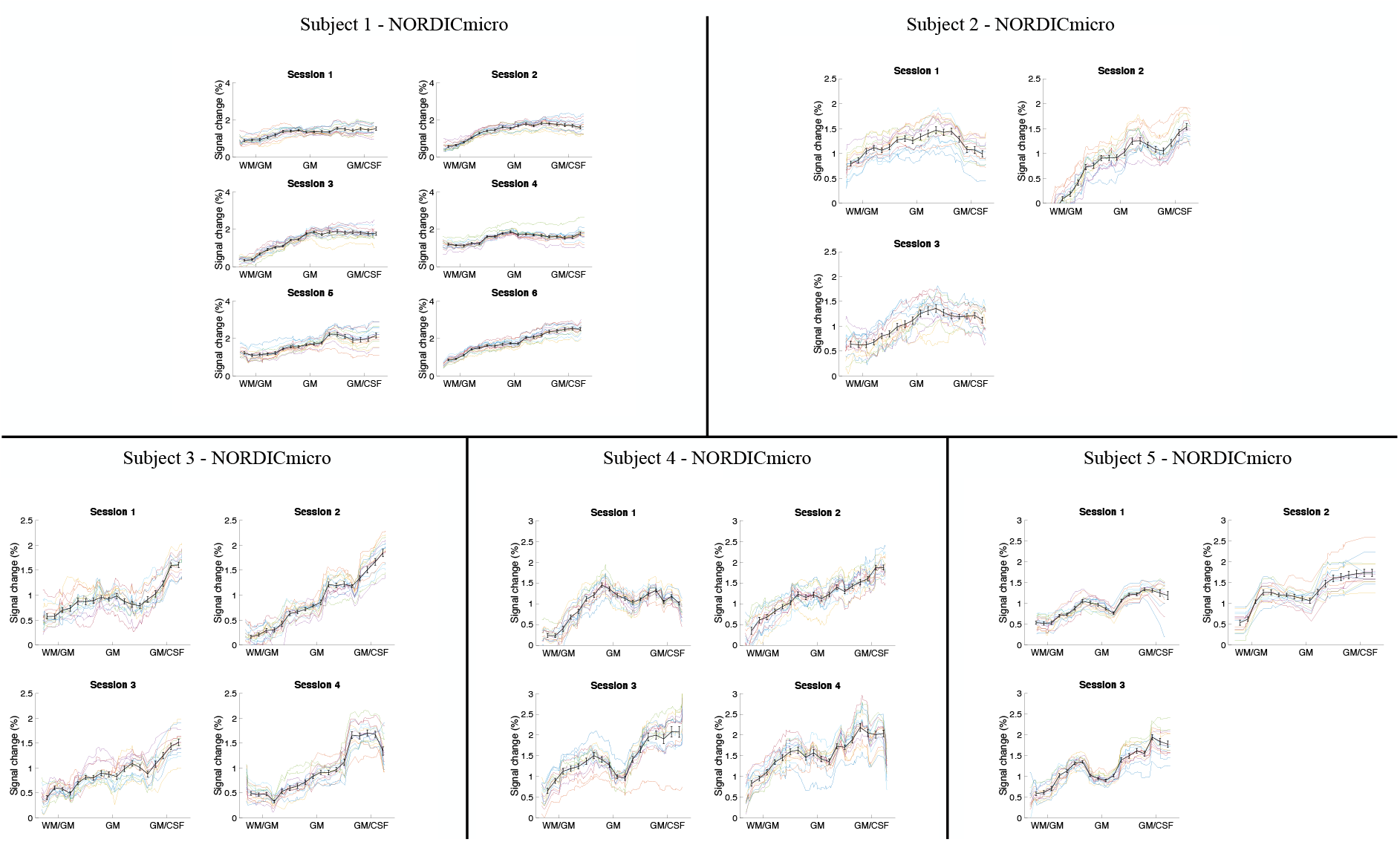
NORDICmicro layerprofiles of all sessions and subjects. Colored profiles represent single trials with across-trial mean profiles plotted on top in black. Individual trial profiles seem to closely follow the mean, implying robust laminar patterns of activation within sessions. Error bars = SEM across trials (N = 15).

**Figure 3.**
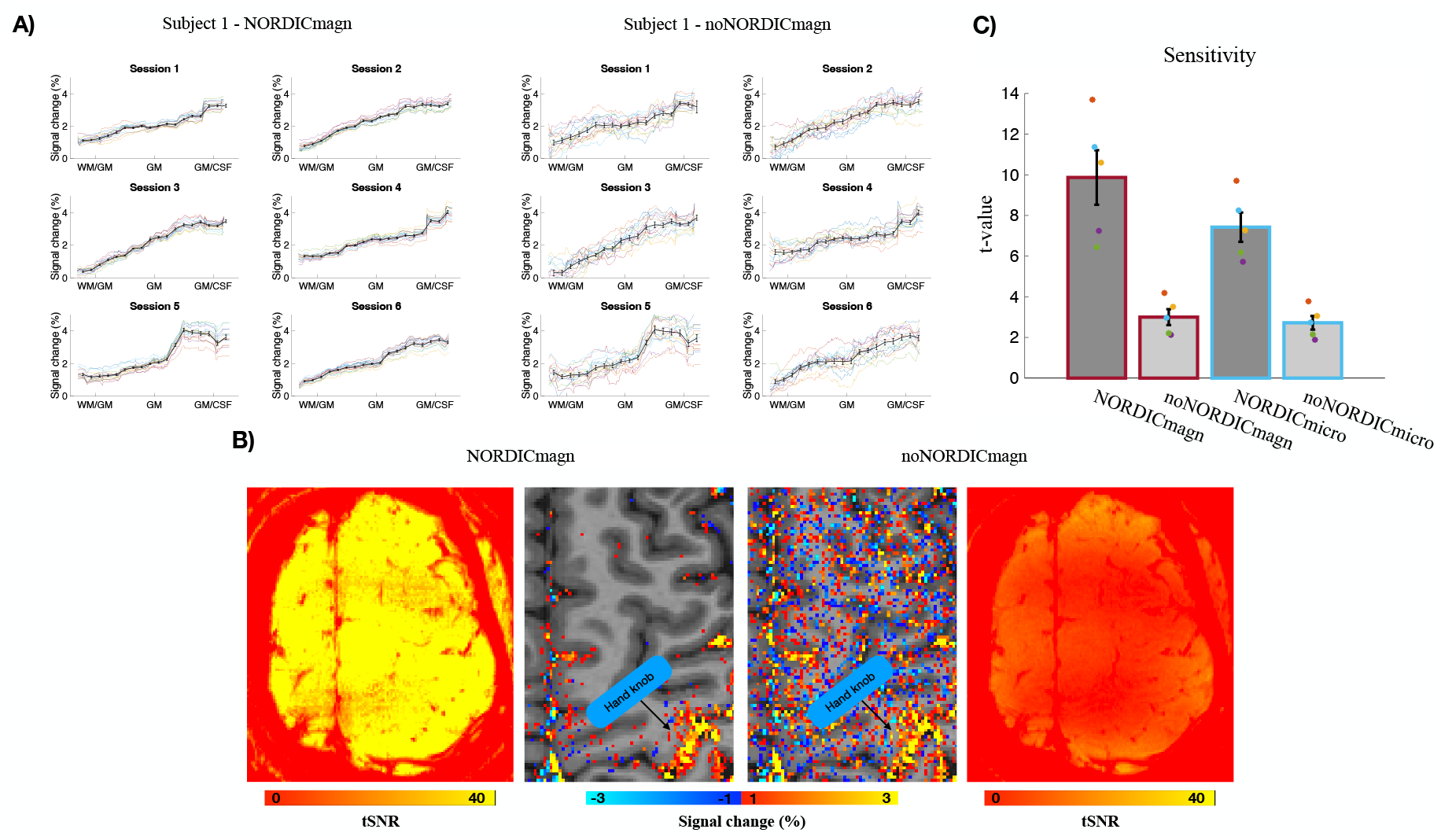
Effect of NORDIC on signal stability ***A)*** NORDICmagn and noNORDICmagn layerprofiles of all sessions from an example subject. Colored profiles represent single trials with across-trial mean profiles plotted on top in black. Mean profiles are practically identical across the two data versions, whereas variability around the mean is notably higher without NORDIC denoising. Error bars represent SEM across trials (N = 15). ***B)*** NORDICmagn and noNORDICmagn tSNR and activation maps in an example slice from the same subject as in *A*. Comparison suggests reduced false positive activation after denoising as a result of improved tSNR. ***C)*** Average within-ROI t-values across subjects for the 4 different data versions. Colored dots represent datapoints of each subject. Error bars represent SEM across subjects (N = 5).

A recent 7T layer-fMRI study reported that phase regression came along with reduced sensitivity as measured by the CNR of observed functional responses (Stanley et al., 2020). To examine for such an effect in the present setup, across-subject mean t-values of the 4 different data versions were computed (Figure 3C) and evaluated with ANOVA. Figure 3C shows that the difference in sensitivity between magnitude- and micro data versions was larger when NORDIC was applied compared to when it was not (*NORDICversion* by *veinCorrection* interaction, p = 0.025). Phase regression is thus accompanied by reduced sensitivity, and interestingly, that effect is larger when NORDIC is applied. However, as will be addressed in later sections, part of the effect is likely explained by reduced sensitivity towards spatially unspecific draining veins and is thus desired. The large difference in t-values between NORDIC and noNORDIC applications, observable in Figure 3C, was statistically significant (main effect of *NORDICversion*, p = 0.001).

Figure 4A displays NORDICmagn and NORDICmicro profiles for each session plotted together for each subject. Robustness across-sessions was estimated for NORDICmicro as the standard deviation over sessions divided by the mean (done separately for each layer and then averaged), which we refer to as session-CV. The average session-CV across subjects was 21.91 % ±5.05 %. Figure 4A further illustrates that the difference in percent signal change between magnitude and micro profiles is on average largest in CSF and decreases progressively towards WM, implying that bias towards superficial layers is reduced in the micro profiles. Figure 4B shows NORDICmagn and NORDICmicro hand knob activation maps in an example slice from the first session of each subject. Voxels with large response suppression after phase regression are predominantly located where large veins are expected, i.e., towards superficial layers and CSF, supporting the notion that phase regression mainly suppresses macrovascular signal contributions. While the majority of voxels likely to contain large veins (i.e., superficial/CSF voxels with very strong activation in magnitude maps) appear to have strongly suppressed responses after phase regression, there is obvious cases (blue arrows points to examples of this) where the method left residual macrovascular signal, highlighting that although the method reduces macrovascular contributions, it is not perfect.

**Figure 4.**
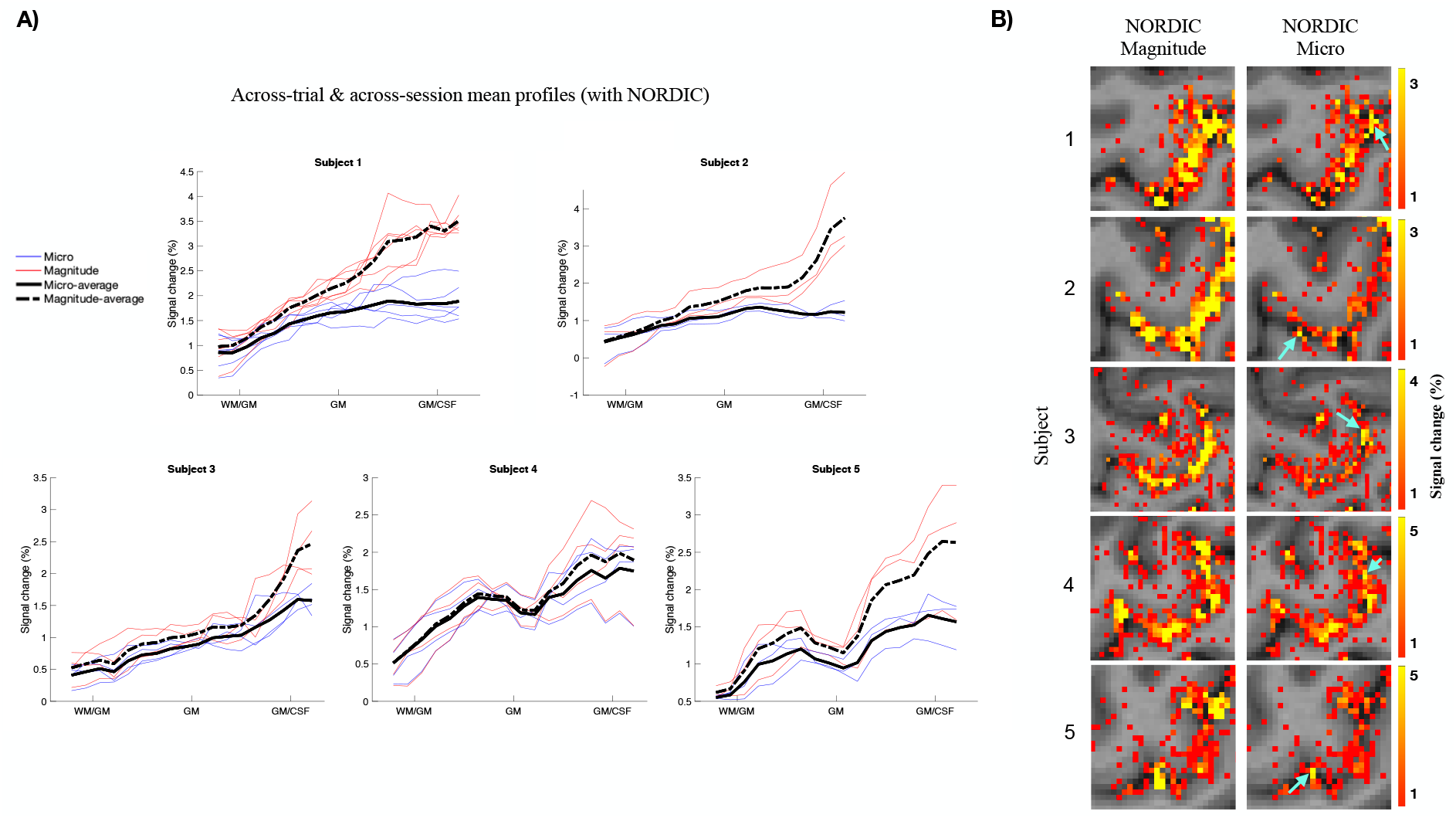
Reproducibility of layer profiles across sessions and the effect of phase regression on superficial bias ***A)*** NORDICmagn and NORDICmicro layerprofiles plotted together for all subjects. Colored profiles represent single sessions (red = magnitude and blue = micro) with across-session mean profiles plotted on top in black (dashed = magnitude and solid = micro). Profiles appear consistent across sessions with seemingly smaller superficial bias after phase regression. ***B)*** NORDICmagn and NORDICmicro hand knob activation maps in example slices from each subject. Although most voxels likely to contain large veins (see main text) have strongly suppressed responses after phase regression, residual macrovascular contributions are still left in NORDICmicro activation maps as pointed out by the blue arrows.

Since the effectiveness of phase regression is CNR-dependent, we suspected smaller superficial bias in NORDICmicro compared to noNORDICmicro profiles. This was confirmed using ANOVA with slopes of linear fits as the dependent variable, which revealed a significant *NORDICversion* by *veinCorrection* interaction (p = 0.026). That is, the difference in slopes between magnitude- and micro data versions was larger when NORDIC was applied compared to when it was not (see average slope across subjects for each data version in Figure 5A). This effect is further visualized in Figure 5B, which illustrates that noNORDICmicro across-session mean profiles of each subject have steeper gradients from WM to CSF compared to NORDICmicro (Bonferroni corrected Post hoc comparison, p = 0.017). To accommodate the potential risk of slopes being biased towards larger values due to our ROI selection criteria, we made an additional version of Figure 5 where linear fits were computed based on superficial depths only which is shown in Figure S3.

**Figure 5.**
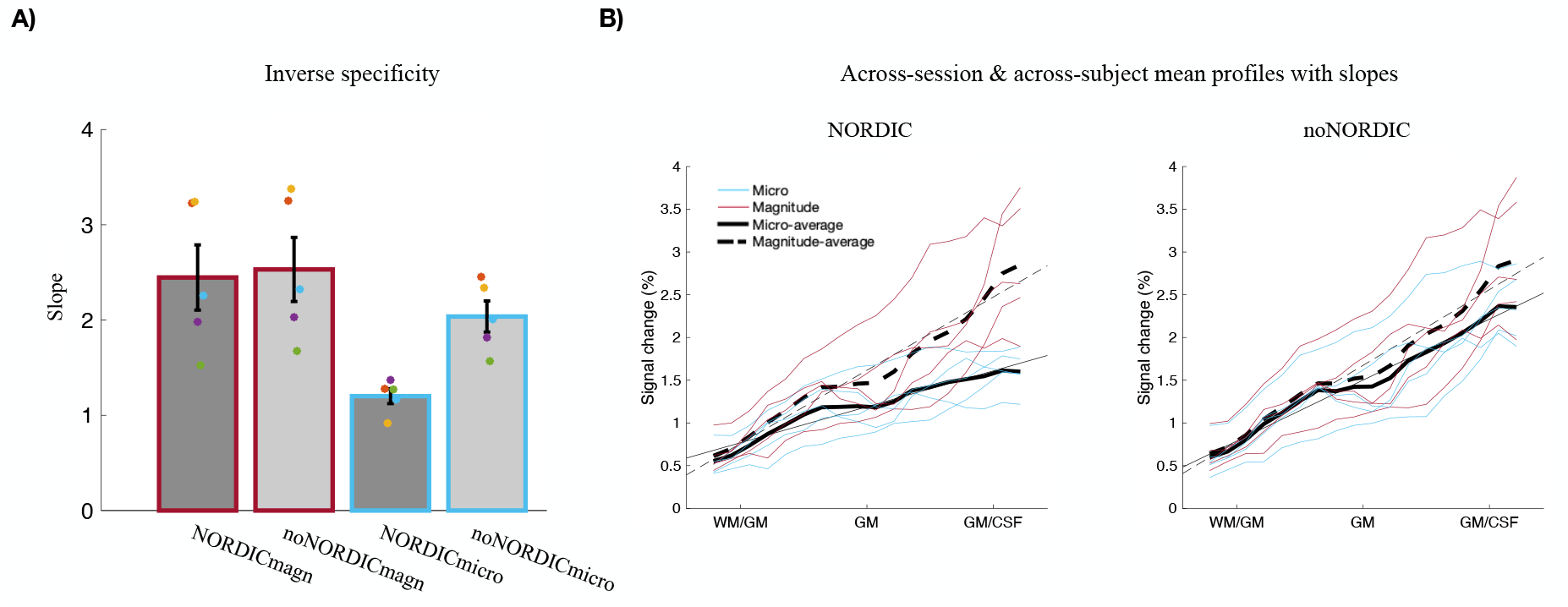
Effect of phase regression on superficial bias ***A)*** Average of slopes of linear fits to each subject’s mean layerprofile for the 4 different data versions (colored profiles in *B)*. Colored dots represent datapoints of each subject. Error bars represent SEM across subjects (N = 5). ***B)*** Magnitude and micro layerprofiles plotted together for all subjects both with and without NORDIC. Colored profiles represent single subjects (red = magnitude and blue = micro) with across-subject mean profiles plotted on top in black (dashed = magnitude and solid = micro). Straight lines indicate linear fits to each of the 4 mean profiles. The difference in slopes between magnitude and micro profiles is significantly larger with NORDIC, suggesting a more effective phase regression compared to without NORDIC.

## 4. Discussion

The aim of the present study was to test the feasibility of L-fMRI at 3T with GE-BOLD by use of advanced post processing. 3T fMRI with submillimeter spatial resolution suffers from poor signal stability which partly explains why nearly all existing L-fMRI studies were performed at 7T systems. The results presented here demonstrate how this challenge can be overcome with NORDIC denoising, allowing for reliable detection of laminar activation both within and across sessions at 3T. This can be achieved with standard MR hardware, without application of explicit spatial smoothing kernels which degrade spatial specificity, and in a single functional run of 18 minutes (including functional localizer) making it highly practical. Additionally, we tested phase regression (Menon, 2002) as a candidate method to reduce superficial bias caused by large veins and found evidence for significant improvements in obtained layer profiles, although some macrovascular contribution likely remained.

### 4.1 Sensitivity and within-session reliability

NORDICmicro profiles are depicted in Figure 2 for each experimental session. At the level of individual trials, profiles seem highly reproducible. This is supported by a small trial-CV of 20.84 % ±3.70 % across subjects for NORDICmicro. CV-estimates for single trials are to our knowledge not reported in the L-fMRI literature, but in Huber, Tse, et al., (2018) the CV across runs was 25 % for both VASO and BOLD. A prerequisite for reproducible single trial estimates is a stable underlying signal which can be quantified as tSNR. Here it was computed from motion corrected and detrended NORDICmagn data before motion and RETROICOR regressors were filtered out to avoid uneven comparison with previous ultrahigh field (≥7T) submillimeter studies. The average tSNR within ROIs was 39.47 ±5.07 across subjects, which is comparable to, if not higher than what is commonly reported for submillimeter 7T GE-BOLD setups (Aitken et al., 2020; Beckett et al., 2020; Huber, Ivanov, et al., 2018; Rua et al., 2017; Stanley et al., 2020; Zaretskaya et al., 2020). Huber, Ivanov, et al., (2018), Rua et al., (2017) and Aitken et al., (2020), for example, reported tSNR value across subjects of 39, ∼20 and 12.5, respectively, in motor and visual cortex ROIs. Note that tSNR depends on factors such as EPI-readout strategy (2D versus 3D acquisitions), acquisition parameters (repetition time, flip angle, etc.), voxel size, and ROI selection (all brain voxels versus specific subsets). Choices in sequence design, preprocessing and ROI selection may thus partly explain how the present 3T setup has comparable tSNR values to 7T observations. However, the main reason is likely the application of NORDIC denoising, which is apparent from the fact that in our sample it increased tSNR by a factor of ∼3 on average. Importantly, NORDIC increases tSNR without adding any noteworthy spatial smoothness to the images (Dowdle et al., 2021; Vizioli et al., 2021). In one of the original NORDIC articles, the developers found that denoising on average increased tSNR by a factor of more than 2 (Vizioli et al., 2021). The larger observed effect of NORDIC in the present data might partly be explained by the fact that methods for estimating the threshold under which components with lower singular values were discarded, differed across studies (see Methods section 2.4.1, Figure S1 and Figure S2). Also, in contrast to Vizioli et al., (2021), we used the version of NORDIC that works in image space as opposed to raw data space, without the need for an appended noise volume and g-factor map. A complementary explanation is that the degree to which thermal noise is dominant, scales inversely with field strength (Triantafyllou et al., 2005). Loosely speaking, this implies that more noise is present to begin with and thus available for removal at 3T.

Accordingly, 3T submillimeter fMRI with NORDIC denoising can achieve within session stability measures comparable to or in some cases higher than what has been previously reported at ultrahigh field strengths. The outcome is sufficient sensitivity to robustly detect task induced BOLD responses which is underlined by the average t-value of ROI voxels being 7.43 ±1.61 across subjects for NORDICmicro (14 degrees of freedom, assuming independent trial beta-estimates) (Figure 3C).

### 4.2 Across-session reliability

Each subject was scanned in multiple sessions on multiple days to assess across-session reproducibility of observed layer profiles. To examine the result, we depicted the mean profiles of all sessions for each subject in the same plot (Figure 4A). Visual assessment indicates that individual session profiles consistently followed the across-session mean profiles, both regarding the laminar pattern and response magnitude. This is quantified as an average session-CV for NORDICmicro of 21.91 % ±5.05 % across subjects, which, like the trial-CV estimate, is comparable to the across-days CV-estimate observed for VASO and BOLD (25 % and 30 %, respectively) in the previously mentioned 9.4T study (Huber, Tse, et al., 2018). For reference, non-laminar supramillimeter studies found CV’s of 24 % and ∼28 % (means across subjects) for the variability of response magnitudes during motor tasks in M1 ROIs across 3 sessions at 3T (Tjandra et al., 2005) and 7T (Krieger et al., 2014), respectively. M1 L-fMRI studies that scanned the same participants in multiple sessions and where associated layerprofiles of individual sessions were reported are sparse making comparison difficult. However, this was done in Chai et al., (2020) who also used a finger tapping paradigm and focused the analysis onto ROIs from the hand knob, although with a so-called VAPER sequence rather than GE-BOLD. No quantitative estimate was presented, but visual comparison with the profiles from that study further supports that an across-session reliability in agreement with previous M1 studies, across field strengths and spatial resolutions, could be achieved despite the challenges of submillimeter fMRI at 3T.

### 4.3 Specificity

GE-EPI is the most frequently applied sequence in layer-fMRI studies due to its superior statistical efficiency and ease of implementation. However, its weighting towards macrovasculature is well established (Menon, 2012; Turner, 2002; Uludağ et al., 2009; Yacoub et al., 2003) and needs to be dealt with for laminar purposes where spatial specificity is of the essence. The superficial bias that follows from large vein sensitivity was clearly present in the NORDICmagn profiles which indeed possessed the characteristic positive gradient from WM to CSF (Figure 4A and Figure 5B) consistently found with GE-BOLD (Aitken et al., 2020; de Hollander et al., 2021; Han et al., 2021; Huber et al., 2015; Kok et al., 2016; Shao et al., 2021; Stanley et al., 2020) (see also Figure S3). Also, activation maps revealed higher percent signal changes towards the surface and in CSF which adds to the evidence for large vein contamination (Figure 4B). Phase regression has recently shown promise as a method to reduce the macrovascular sensitivity of GE-BOLD in 7T submillimeter settings to obtain microvascular specificity comparable to regular SE-BOLD (Stanley et al., 2020, but see Han et al., 2021 for SE-sequence with improved microvascular weighting), and we thus decided to test its effectiveness in the present 3T setting. If phase regression predominantly reduces signal originating from large veins, we should see larger signal suppression towards the surface and in CSF. We did indeed find evidence for this in line with previous observations (Curtis et al., 2014; Menon, 2002; Stanley et al., 2020; Vicente et al., 2021); the difference in magnitudes between NORDICmagn and NORDICmicro profiles progressively increased from WM to CSF (Figure 4A) and the same effect is observable directly in activation maps (Figure 4B). The degree of superficial bias was estimated as the slope of the linear fit to across-session mean profiles of each subject inspired by previous efforts (Beckett et al., 2020; De Martino et al., 2013; Huber et al., 2017) and the extent of the effect is then reflected in the decrease of the slope from NORDICmagn to NORDICmicro which is shown in Figure 5A. Interestingly, the effect of phase regression, as measured by its impact on slopes, was significantly larger when NORDIC was applied (Figure 5A and Figure 5B), illustrating its dependency on CNR of both the magnitude and phase time series. This is supported by a significantly larger reduction in t-values after phase regression for NORDIC compared to noNORDIC (Figure 3C), which we interpret to mainly reflect a reduced sensitivity towards macrovasculature. Although the slope might be a simplified measure of inverse specificity, as also pointed out by Beckett et al., (2020), it does illustrate that phase regression substantially reduced the gradient from WM to CSF which is generally believed to originate from large veins.

That being said, several lines of evidence indicate that non-negligible contributions from macrovascular sources were still in play. First, even though the group layer-profile for NORDICmicro tends to flatten towards CSF as compared to the NORDICmagn profile which keeps rising, the peak is located at the GM/CSF border, whereas sequences proven to have higher spatial specificity peaks within the GM of M1 during finger tapping (VASO/VAPER: (Beckett et al., 2020; Chai et al., 2020; Huber et al., 2017; Persichetti et al., 2020), SE/GRASE: (Beckett et al., 2020; Han et al., 2021), ASL: (Shao et al., 2021)). A peak towards CSF after phase regression was also observed for the visual cortex in Stanley et al., (2020), whereas neural activity is expected to peak in middle layers during visual stimulation (Liu et al., 2020). Second, the lateral end of the hand knob (BA4a) in M1 has typically been associated with a double peak response when examined with more spatially specific sequences at 7T (Beckett et al., 2020; Chai et al., 2020; Guidi et al., 2020; Huber et al., 2017; Shao et al., 2021). This feature has been used as a hallmark for spatial specificity and is accordingly less robustly observed with GE-BOLD (Huber et al., 2017) where peaks become less distinct as a result of signal drainage from deep layers to the surface. While NORDICmicro profiles from 2 of 5 subjects in the present experiment consistently had the double peak feature (Figure 2), the group profile can better be described by a superficial peak with a shoulder in the deep layers (Figure 5B). Although we do believe that the lack of two distinct peaks is partly explained by macrovascular influence, it may also be explained by the fact that we extract signal from relatively large ROIs spanning several slices (Han et al., 2021; Pais-Roldán et al., 2020). Moreover, not all subjects have double peak profiles even with non-GE-BOLD sequences (Beckett et al., 2020; Shao et al., 2021), thus we cannot rule out that more distinct peaks would emerge in the group profile with different subjects. Third, NORDICmicro activation maps still contain CSF voxels with strong percent signal changes (Figure 4B). Finally, it is not completely clear how well the microvascular signal is preserved in GM voxels which partially volume with a large vein. To properly separate micro and macrovascular signal components the method assumes temporal uncoupling between hemodynamic responses of the two compartments (Stanley et al., 2020). Considering that temporal delays between the two types of vasculatures are on the order of about 1-3 seconds (Kay et al., 2020), it may be too subtle to be resolved with TRs commonly employed within the field.

In summary, the results presented here support that phase regression substantially reduces superficial bias and the associated decrease in sensitivity likely reflects a desired feature of reduced macrovascular weighting. It comes, however, with some imperfections that need to be considered when selecting a tool to handle the problem of veins which is mandatory in virtually any L-fMRI application.

### 4.4 Previous 3T submillimeter studies

The present demonstration of L-fMRI being feasible at 3T with GE-BOLD is not the first; several previous studies have successfully obtained layer-dependent measures of activation (Koopmans et al., 2010; Markuerkiaga et al., 2020; Puckett et al., 2016; Ress et al., 2007; Scheeringa et al., 2016). However, these studies relied on acquisition and analysis strategies which may not be generally applicable. For example, Markuerkiaga et al., (2020) and Koopmans et al., (2010) used FLASH-sequences to measure BOLD responses with TRs of 130 s and 60 s, respectively. In Ress et al., (2007) specialized hardware was used to overcome SNR issues, which was solved in Puckett et al., (2016) by using long scan durations (18-25 3-minute runs per subject), and in Scheeringa et al., (2016) by only analyzing vertices with the 10 % highest t-values and by averaging over large areas. In addition to BOLD, exciting results from two recent conference abstracts by Laurentius Huber and colleagues indicate that CBV-weighted fMRI with SS-SI-VASO might be a viable tool for measuring layer-dependent responses at 3T with excellent spatial specificity although with a tradeoff of reduced CNR (L. Huber, Kronbichler, Stirnberg, Kronbichler, et al., 2021; L. Huber, Kronbichler, Stirnberg, Poser, et al., 2021). The present results add to the contributions of these studies by showing that with the implementation of NORDIC, reliable submillimeter activation maps can be obtained with very high statistical efficiency and without the need for long scan durations, extensive averaging, specialized hardware etc.

### 4.5 Choice of NORDIC version

Changing the FE argument to 1.15 (default = 1) in NORDIC, as done here, effectively increases the number of removed time series components. This leads to a dramatic increase in tSNR, but does come with the risk of removing signals of interest. For the present setup, removed components appeared to be dominated by thermal noise (Figure S1) and although we did find a significant reduction in the magnitude of group-averaged layer profiles, the effect was small compared to error associated with estimation of group-mean responses (Figure S2). For this reason, and given the CNR dependency related to the effectiveness of phase regression, we found it optimal to adjust this NORDIC parameter in the present case. However, the magnitude of the effect likely depends on sequence, acquisition parameters, field strength, etc. Alteration of the parameter should thus be done with care and might not even be necessary considering that the average tSNR across sessions and subjects was 22.58 ±2.66 when NORDIC was applied with default settings (Figure S1).

### 4.6 Limitations and future work

We opted to prioritize multiple test-retest scans on the same subjects to examine reliability of the current 3T setup. However, we acknowledge that the small sample size of only 5 subjects and 20 total sessions precludes rigorous statistical analysis and the reported reliability measures, as well as the comparison with such estimates from e.g. Huber, Tse, et al., (2018) which were also based on small samples, should be interpreted in that light. Future test-retest studies with larger sample sizes would thus be beneficial to further test our suggestion that layer-dependent activation can be reliably measured at 3T. Such studies are to our knowledge currently lacking in the field as a whole, including ultrahigh field, and would be valuable as the validity of any method ultimately depends on its reliability. Moreover, as discussed above, the 3T GE-BOLD setup presented here is associated with high sensitivity but limited spatial specificity which is also true for GE-BOLD at higher field strengths. Given that every L-fMRI study has vastly different requirements regarding sensitivity, specificity, temporal resolution etc., future work at 3T could ideally test alternative sequences known to excel in areas where GE-BOLD is challenged. An example of a candidate sequence is SS-SI-VASO which already showed great promise at 3T (L. Huber, Kronbichler, Stirnberg, Kronbichler, et al., 2021; L. Huber, Kronbichler, Stirnberg, Poser, et al., 2021) and might be desired over GE-BOLD when specificity is prioritized over sensitivity. The focus could, along this line, be directed towards alternative postprocessing methods for deveining, with TDM (Kay et al., 2020) being an exciting suitor.

Finally, it should be mentioned that certain compromises were necessary in order to achieve submillimeter resolution functional images with a reasonable temporal resolution: 1) The FOV was small (26 axial slices of 0.82 mm thickness) which in some cases led to fold-over artifacts visible in the magnitude images. Although this did not seem to have a big effect on the time series data in the present study, it should be of priority to mitigate the artifact by making sure the imaging slab extends outside the brain which comes at the cost of reduced brain coverage. Alternatively, the FOV can be increased in a tradeoff with longer TRs; 2) Partial Fourier reconstruction was employed for faster coverage of K-space but comes with a reduction in the effective spatial resolution and it might introduce image artifacts in locations of high frequency phase changes (Haacke et al., 1991). Partial Fourier was implemented with the zero-filling approach in line with previous studies using NORDIC (Vizioli et al., 2021) and phase-regression (Stanley et al., 2020). More advanced methods such as POCS (Haacke et al., 1991) might be beneficiary in order to minimize spatial blurring. Alternatively, Partial Fourier could be omitted entirely at the cost of temporal resolution. In future studies, advanced parallel imaging techniques such as 3D CAIPIRINHA acceleration and multiband excitation can be implemented to increase the coverage and temporal resolution of submillimeter fMRI at 3 T.

## 5. Conclusion

We used GE-BOLD fMRI to test the feasibility of L-fMRI at 3T with NORDIC and phase regression. The results suggest that SNR limitations usually accompanied with submillimeter isotropic voxels at 3T can be overcome with NORDIC denoising. As a result, layer-dependent BOLD responses could be detected reliably within and across sessions with high statistical efficiency. Importantly, this was achievable with standard MR hardware, without application of explicit spatial smoothing kernels, and in a single functional run, making it highly practical. As expected, we observed a strong superficial bias in magnitude layer profiles which was substantially reduced after phase regression, although some macrovascular contributions remained. We believe the present 3T results, coupled with the vast amount of research going into improving microvascular weighting, will help making L-fMRI available to a much wider community.

## Acknowledgements

The authors would like to thank all subjects for their participation, Renzo Huber and Rüdiger Stirnberg for sharing the 3D-EPI sequence, and Steen Müller for support with NORDIC. This work is funded by grants from Sino-Danish Center (SDC).

## Credit authorship contribution statement

**Lasse Knudsen** Conceptualization, Methodology, Formal analysis, Investigation, Writing - original draft, Writing - review and editing, Visualization.

**Christopher Bailey** Conceptualization, Writing -review and editing, Supervision.

**Jakob U. Blicher** Conceptualization, Methodology, Resources, Writing - review and editing, Supervision, Funding acquisition.

**Yan Yang** Conceptualization, Writing - review and editing, Supervision, Project administration

**Peng Zhang** Methodology, Software, Supervision, Writing – review and editing.

**Torben E. Lund** Conceptualization, Methodology, Software, Investigation, Writing - Review & Editing, Supervision.

## References

Aitken, F., Menelaou, G., Warrington, O., Koolschijn, R. S., Corbin, N., Callaghan, M. F., & Kok, P. (2020). Prior expectations evoke stimulus-specific activity in the deep layers of the primary visual cortex. PLoS Biology, 18(12), 1–19. https://doi.org/10.1371/journal.pbio.3001023

Avants, B. B., Tustison, N. J., Song, G., Cook, P. A., Klein, A., & Gee, J. C. (2011). A reproducible evaluation of ANTs similarity metric performance in brain image registration. NeuroImage, 54(3), 2033–2044. https://doi.org/10.1016/j.neuroimage.2010.09.025

Báez-Yánez, M. G., Ehses, P., Mirkes, C., Tsai, P. S., Kleinfeld, D., & Scheffler, K. (2017). The impact of vessel size, orientation and intravascular contribution on the neurovascular fingerprint of BOLD bSSFP fMRI. NeuroImage, 163(August), 13–23. https://doi.org/10.1016/j.neuroimage.2017.09.015

Bandettini, P. A., Huber, L., & Finn, E. S. (2021). Challenges and opportunities of mesoscopic brain mapping with fMRI. Current Opinion in Behavioral Sciences, 40, 189–200. https://doi.org/10.1016/j.cobeha.2021.06.002

Beckett, A. J. S., Dadakova, T., Townsend, J., Huber, L., Park, S., & Feinberg, D. A. (2020). Comparison of BOLD and CBV using 3D EPI and 3D GRASE for cortical layer functional MRI at 7 T. Magnetic Resonance in Medicine, 84(6), 3128–3145. https://doi.org/10.1002/mrm.28347

Chai, Y., Li, L., Huber, L., Poser, B. A., & Bandettini, P. A. (2020). Integrated VASO and perfusion contrast: A new tool for laminar functional MRI. NeuroImage, 207(August 2019), 116358. https://doi.org/10.1016/j.neuroimage.2019.116358

Chaimow, D., Yacoub, E., Uğurbil, K., & Shmuel, A. (2018). Spatial specificity of the functional MRI blood oxygenation response relative to neuronal activity. NeuroImage, 164(August), 32–47. https://doi.org/10.1016/j.neuroimage.2017.08.077

Cox, R. W. (1996). AFNI: Software for analysis and visualization of functional magnetic resonance neuroimages. Computers and Biomedical Research, 29(3), 162–173. https://doi.org/10.1006/cbmr.1996.0014

Curtis, A. T., Hutchison, R. M., & Menon, R. S. (2014). Phase based venous suppression in resting-state BOLD GE-fMRI. NeuroImage, 100, 51–59. https://doi.org/10.1016/j.neuroimage.2014.05.079

de Hollander, G., van der Zwaag, W., Qian, C., Zhang, P., & Knapen, T. (2021). Ultra-high field fMRI reveals origins of feedforward and feedback activity within laminae of human ocular dominance columns. NeuroImage, 228, 117683. https://doi.org/10.1016/j.neuroimage.2020.117683

De Martino, F., Moerel, M., Ugurbil, K., Goebel, R., Yacoub, E., & Formisano, E. (2015). Frequency preference and attention effects across cortical depths in the human primary auditory cortex. Proceedings of the National Academy of Sciences, 112(52), 16036–16041. https://doi.org/10.1073/pnas.1507552112

De Martino, F., Zimmermann, J., Muckli, L., Ugurbil, K., Yacoub, E., & Goebel, R. (2013). Cortical Depth Dependent Functional Responses in Humans at 7T: Improved Specificity with 3D GRASE. PLoS ONE, 8(3), 30–32. https://doi.org/10.1371/journal.pone.0060514

Dowdle, L. T., Vizioli, L., Moeller, S., Olman, C., Ghose, G., Yacoub, E., & Ugurbil, K. (2021). Improving Sensitivity to Functional Responses without a Loss of Spatiotemporal Precision in Human Brain Imaging. BioRxiv, 2021.08.26.457833. https://www.biorxiv.org/content/10.1101/2021.08.26.457833v1%0Ahttps://www.biorxiv.org/content/10.1101/2021.08.26.457833v1.abstract

Dumoulin, S. O., Fracasso, A., van der Zwaag, W., Siero, J. C. W., & Petridou, N. (2018). Ultra-high field MRI: Advancing systems neuroscience towards mesoscopic human brain function. NeuroImage, 168(September), 345–357. https://doi.org/10.1016/j.neuroimage.2017.01.028

Duvernoy, H. M., Delon, S., & Vannson, J. L. (1981). Cortical blood vessels of the human brain. Brain Research Bulletin, 7(5), 519–579. https://doi.org/10.1016/0361-9230(81)90007-1

Felleman, D. J., & Van Essen, D. C. (1991). Distributed Hierarchical Processing in the Primate Cerebral Cortex. Cerebral Cortex, 1(1), 1–47. https://doi.org/10.1093/cercor/1.1.1

Friston, K. J., Williams, S., Howard, R., Frackowiak, R. S. J., & Turner, R. (1996). Movement-Related effects in fMRI time-series. Magnetic Resonance in Medicine, 35(3), 346–355. https://doi.org/10.1002/mrm.1910350312

Glover, G. H., Li, T.-Q., & Ress, D. (2000). Image-based method for retrospective correction of physiological motion effects in fMRI: RETROICOR. Magnetic Resonance in Medicine, 44(1), 162–167. https://doi.org/10.1002/1522-2594(200007)44:1<162::AID-MRM23>3.0.CO;2-E

Goense, J. B. M., & Logothetis, N. K. (2006). Laminar specificity in monkey V1 using high-resolution SE-fMRI. Magnetic Resonance Imaging, 24(4), 381–392. https://doi.org/10.1016/j.mri.2005.12.032

Guidi, M., Huber, L., Lampe, L., Merola, A., Ihle, K., & Möller, H. E. (2020). Cortical laminar resting-state signal fluctuations scale with the hypercapnic blood oxygenation level-dependent response. Human Brain Mapping, 41(8), 2014–2027. https://doi.org/10.1002/hbm.24926

Haacke, E. M., Lindskogj, E. D., & Lin, W. (1991). A fast, iterative, partial-fourier technique capable of local phase recovery. Journal of Magnetic Resonance (1969), 92(1), 126– 145. https://doi.org/10.1016/0022-2364(91)90253-P

Hall, J. (2022). Linear Deming Regression (https://www.mathworks.com/matlabcentral/fileexchange/33484-linear-deming-regression), zMATLAB Central File Exchange. Retrieved March 3, 2022.

Han, S., Eun, S., Cho, H., Uluda, K., & Kim, S. (2021). NeuroImage Improvement of sensitivity and specificity for laminar BOLD fMRI with double spin-echo EPI in humans at 7 T. 241(July). https://doi.org/10.1016/j.neuroimage.2021.118435

Huang, P., Correia, M. M., Rua, C., Rodgers, C. T., Henson, R. N., & Carlin, J. D. (2021). Correcting for Superficial Bias in 7T Gradient Echo fMRI. Frontiers in Neuroscience, 15(September), 1–16. https://doi.org/10.3389/fnins.2021.715549

Huber, L., Kronbichler, L., Stirnberg, R., Poser, B. A., Fernández-Cabello, S., Stöcker, T., & Kronbichler, M. (2021). Evaluating the capabilities and challenges of layer-fMRI VASO at 3T. Annual Meeting of the Organization for Human Brain Mapping, 1229.

Huber, L., Kronbichler, M., Stirnberg, R., Kronbichler, L., Fernandez-Cabello, S., Stöcker, T., & Poser, B. A. (2021). Does the field strength matter? Comparing layer-fMRI VASO across 3 T, 7 T, and 9.4 T. ESMRMB 2021, 34, 1–204. https://doi.org/10.1007/s10334-021-00947-8

Huber, Laurentius, Goense, J., Kennerley, A. J., Trampel, R., Guidi, M., Reimer, E., Ivanov, D., Neef, N., Gauthier, C. J., Turner, R., & Möller, H. E. (2015). Cortical lamina-dependent blood volume changes in human brain at 7T. NeuroImage, 107, 23–33. https://doi.org/10.1016/j.neuroimage.2014.11.046

Huber, Laurentius, Handwerker, D. A., Jangraw, D. C., Chen, G., Hall, A., Stüber, C., Gonzalez-Castillo, J., Ivanov, D., Marrett, S., Guidi, M., Goense, J., Poser, B. A., & Bandettini, P. A. (2017). High-Resolution CBV-fMRI Allows Mapping of Laminar Activity and Connectivity of Cortical Input and Output in Human M1. Neuron, 96(6), 1253-1263.e7. https://doi.org/10.1016/j.neuron.2017.11.005

Huber, Laurentius, Ivanov, D., Handwerker, D. A., Marrett, S., Guidi, M., Uludağ, K., Bandettini, P. A., & Poser, B. A. (2018). Techniques for blood volume fMRI with VASO: From low-resolution mapping towards sub-millimeter layer-dependent applications. NeuroImage, 164(November), 131–143. https://doi.org/10.1016/j.neuroimage.2016.11.039

Huber, Laurentius, Ivanov, D., Krieger, S. N., Streicher, M. N., Turner, R., Mildner, T., Poser, B. A., & Harald, E. M. (2014). Slab-Selective, BOLD-Corrected VASO at 7 Tesla Provides Measures of Cerebral Blood Volume Reactivity with High Signal-to-Noise Ratio. 148, 137–148. https://doi.org/10.1002/mrm.24916

Huber, Laurentius, Poser, B. A., Bandettini, P. A., Arora, K., Wagstyl, K., Cho, S., Goense, J., Nothnagel, N., Morgan, A. T., van den Hurk, J., Müller, A. K., Reynolds, R. C., Glen, D. R., Goebel, R., & Gulban, O. F. (2021). LayNii: A software suite for layer-fMRI. NeuroImage, 237(July), 118091. https://doi.org/10.1016/j.neuroimage.2021.118091

Huber, Laurentius, Tse, D. H. Y., Wiggins, C. J., Uludağ, K., Kashyap, S., Jangraw, D. C., Bandettini, P. A., Poser, B. A., & Ivanov, D. (2018). Ultra-high resolution blood volume fMRI and BOLD fMRI in humans at 9.4 T: Capabilities and challenges. NeuroImage, 178(May), 769–779. https://doi.org/10.1016/j.neuroimage.2018.06.025

Jochimsen, T. H., Norris, D. G., Mildner, T., & Möller, H. E. (2004). Quantifying the intra-and extravascular contributions to spin-echo fMRI at 3 T. Magnetic Resonance in Medicine, 52(4), 724–732. https://doi.org/10.1002/mrm.20221

Kay, K., Jamison, K. W., Vizioli, L., Zhang, R., Margalit, E., & Ugurbil, K. (2019). A critical assessment of data quality and venous effects in sub-millimeter fMRI. NeuroImage, 189, 847–869. https://doi.org/10.1016/j.neuroimage.2019.02.006

Kay, K., Jamison, K. W., Zhang, R. Y., & Uğurbil, K. (2020). A temporal decomposition method for identifying venous effects in task-based fMRI. Nature Methods, 17(10), 1033–1039. https://doi.org/10.1038/s41592-020-0941-6

Kok, P., Bains, L. J., Van Mourik, T., Norris, D. G., & De Lange, F. P. (2016). Selective activation of the deep layers of the human primary visual cortex by top-down feedback. Current Biology, 26(3), 371–376. https://doi.org/10.1016/j.cub.2015.12.038

Koopmans, P. J., Barth, M., & Norris, D. G. (2010). Layer-specific BOLD activation in human V1. Human Brain Mapping, 31(9), 1297–1304. https://doi.org/10.1002/hbm.20936

Koopmans, P. J., & Yacoub, E. (2019). Strategies and prospects for cortical depth dependent T2 and T2* weighted BOLD fMRI studies. NeuroImage, 197(February), 668–676. https://doi.org/10.1016/j.neuroimage.2019.03.024

Krieger, S. N., Gauthier, C. J., Ivanov, D., Huber, L., Roggenhofer, E., Sehm, B., Turner, R., & Egan, G. F. (2014). Regional reproducibility of calibrated BOLD functional MRI: Implications for the study of cognition and plasticity. NeuroImage, 101, 8–20. https://doi.org/10.1016/j.neuroimage.2014.06.072

Liu, C., Guo, F., Qian, C., Zhang, Z., Sun, K., Wang, D. J., He, S., & Zhang, P. (2020). Layer-dependent multiplicative effects of spatial attention on contrast responses in human early visual cortex. Progress in Neurobiology, 101897. https://doi.org/10.1016/j.pneurobio.2020.101897

Lu, H., Golay, X., Pekar, J. J., & Van Zijl, P. C. M. (2003). Functional magnetic resonance imaging based on changes in vascular space occupancy. Magnetic Resonance in Medicine, 50(2), 263–274. https://doi.org/10.1002/mrm.10519

Markuerkiaga, I., Marques, J., Bains, L., & Norris, D. (2020). An in-vivo study of BOLD laminar responses as a function of echo time and static magnetic field strength. 1–28. https://doi.org/10.1101/2020.07.16.206383

Marques, J. P., Kober, T., Krueger, G., van der Zwaag, W., Van de Moortele, P.-F., & Gruetter, R. (2010). MP2RAGE, a self bias-field corrected sequence for improved segmentation and T1-mapping at high field. NeuroImage, 49(2), 1271–1281. https://doi.org/10.1016/j.neuroimage.2009.10.002

McColgan, P., Joubert, J., Tabrizi, S. J., & Rees, G. (2020). The human motor cortex microcircuit: insights for neurodegenerative disease. Nature Reviews Neuroscience, 21(8), 401–415. https://doi.org/10.1038/s41583-020-0315-1

Menon, R. S. (2002). Postacquisition suppression of large-vessel BOLD signals in high-resolution fMRI. Magnetic Resonance in Medicine, 47(1), 1–9. https://doi.org/10.1002/mrm.10041

Menon, R. S. (2012). The great brain versus vein debate. NeuroImage, 62(2), 970–974. https://doi.org/10.1016/j.neuroimage.2011.09.005

Moeller, S., Pisharady, P. K., Ramanna, S., Lenglet, C., Wu, X., Dowdle, L., Yacoub, E., Uğurbil, K., & Akçakaya, M. (2021). NOise reduction with DIstribution Corrected (NORDIC) PCA in dMRI with complex-valued parameter-free locally low-rank processing. NeuroImage, 226(October 2020). https://doi.org/10.1016/j.neuroimage.2020.117539

Moerel, M., De Martino, F., Uğurbil, K., Yacoub, E., & Formisano, E. (2019). Processing complexity increases in superficial layers of human primary auditory cortex. Scientific Reports, 9(1), 1–9. https://doi.org/10.1038/s41598-019-41965-w

Muckli, L., De Martino, F., Vizioli, L., Petro, L. S., Smith, F. W., Ugurbil, K., Goebel, R., & Yacoub, E. (2015). Contextual Feedback to Superficial Layers of V1. Current Biology, 25(20), 2690–2695. https://doi.org/10.1016/j.cub.2015.08.057

Pais-Roldán, P., Yun, S. D., Palomero-Gallagher, N., & Shah, N. J. (2020). Cortical depth-dependent human fMRI of resting-state networks using EPIK. BioRxiv, 1–25.

Persichetti, A. S., Avery, J. A., Huber, L., Merriam, E. P., & Martin, A. (2020). Layer-Specific Contributions to Imagined and Executed Hand Movements in Human Primary Motor Cortex. Current Biology, 30(9), 1721-1725.e3. https://doi.org/10.1016/j.cub.2020.02.046

Polimeni, J. R., Renvall, V., Zaretskaya, N., & Fischl, B. (2018). Analysis strategies for high-resolution UHF-fMRI data. NeuroImage, 168, 296–320. https://doi.org/10.1016/j.neuroimage.2017.04.053

Puckett, A. M., Aquino, K. M., Robinson, P. A., Breakspear, M., & Schira, M. M. (2016). The spatiotemporal hemodynamic response function for depth-dependent functional imaging of human cortex. NeuroImage, 139, 240–248. https://doi.org/10.1016/j.neuroimage.2016.06.019

Ress, D., Glover, G. H., Liu, J., & Wandell, B. (2007). Laminar profiles of functional activity in the human brain. NeuroImage, 34(1), 74–84. https://doi.org/10.1016/j.neuroimage.2006.08.020

Rua, C., Costagli, M., Symms, M. R., Biagi, L., Donatelli, G., Cosottini, M., Del Guerra, A., & Tosetti, M. (2017). Characterization of high-resolution Gradient Echo and Spin Echo EPI for fMRI in the human visual cortex at 7 T. Magnetic Resonance Imaging, 40, 98–108. https://doi.org/10.1016/j.mri.2017.04.008

Scheeringa, R., Koopmans, P. J., Van Mourik, T., Jensen, O., & Norris, D. G. (2016). The relationship between oscillatory EEG activity and the laminar-specific BOLD signal. Proceedings of the National Academy of Sciences of the United States of America, 113(24), 6761–6766. https://doi.org/10.1073/pnas.1522577113

Shao, X., Guo, F., Shou, Q., Wang, K., Jann, K., Yan, L., Toga, A. W., Zhang, P., & Wang, D. J. J. (2021). Laminar perfusion imaging with zoomed arterial spin labeling at 7 Tesla. NeuroImage, 245, 118724. https://doi.org/10.1016/j.neuroimage.2021.118724

Sharoh, D., van Mourik, T., Bains, L. J., Segaert, K., Weber, K., Hagoort, P., & Norris, D. G. (2019). Laminar specific fMRI reveals directed interactions in distributed networks during language processing. Proceedings of the National Academy of Sciences of the United States of America, 116(42), 21185–21190. https://doi.org/10.1073/pnas.1907858116

Stanley, O. W., Kuurstra, A. B., Klassen, L. M., Menon, R. S., & Gati, J. S. (2020). Effects of Phase Regression on High-Resolution Functional MRI of the Primary Visual Cortex. NeuroImage, 117631. https://doi.org/10.1016/j.neuroimage.2020.117631

Stirnberg, R., & Stöcker, T. (2021). Segmented K-space blipped-controlled aliasing in parallel imaging for high spatiotemporal resolution EPI. Magnetic Resonance in Medicine, 85(3), 1540–1551. https://doi.org/10.1002/mrm.28486

Tjandra, T., Brooks, J. C. W., Figueiredo, P., Wise, R., Matthews, P. M., & Tracey, I. (2005). Quantitative assessment of the reproducibility of functional activation measured with BOLD and MR perfusion imaging: Implications for clinical trial design. NeuroImage, 27(2), 393–401. https://doi.org/10.1016/j.neuroimage.2005.04.021

Triantafyllou, C., Hoge, R. D., Krueger, G., Wiggins, C. J., Potthast, A., Wiggins, G. C., & Wald, L. L. (2005). Comparison of physiological noise at 1.5 T, 3 T and 7 T and optimization of fMRI acquisition parameters. NeuroImage, 26(1), 243–250. https://doi.org/10.1016/j.neuroimage.2005.01.007

Turner, R. (2002). How much cortex can a vein drain? Downstream dilution of activation-related cerebral blood oxygenation changes. NeuroImage, 16(4), 1062–1067. https://doi.org/10.1006/nimg.2002.1082

Ugurbil, K. (2014). Magnetic Resonance Imaging at Ultrahigh Fields. IEEE Transactions on Biomedical Engineering, 61(5), 1364–1379. https://doi.org/10.1109/TBME.2014.2313619

Uğurbil, K. (2021). Ultrahigh field and ultrahigh resolution fMRI. Current Opinion in Biomedical Engineering, 18, 100288. https://doi.org/10.1016/j.cobme.2021.100288

Uludağ, K., Müller-Bierl, B., & Uğurbil, K. (2009). An integrative model for neuronal activity-induced signal changes for gradient and spin echo functional imaging. NeuroImage, 48(1), 150–165. https://doi.org/10.1016/j.neuroimage.2009.05.051

Vicente, I. De, Uruñuela, E., Termenon, M., & Caballero-gaudes, C. (2021). Dephasing the speaking brain : Cleaning covert sentence production activation maps with a phase-based fMRI data analysis 2. Menon, R. S. (2002). Postacquisition suppression of large - vessel BOLD signals in high - resolution fMRI. Magnetic Resonanc. 2019–2022.

Vizioli, L., Moeller, S., Dowdle, L., Akçakaya, M., De Martino, F., Yacoub, E., & Uğurbil, K. (2021). Lowering the thermal noise barrier in functional brain mapping with magnetic resonance imaging. Nature Communications, 12(1). https://doi.org/10.1038/s41467-021-25431-8

Waehnert, M. D., Dinse, J., Weiss, M., Streicher, M. N., Waehnert, P., Geyer, S., Turner, R., & Bazin, P. L. (2014). Anatomically motivated modeling of cortical laminae. NeuroImage, 93(December), 210–220. https://doi.org/10.1016/j.neuroimage.2013.03.078

Walsh, D. O., Gmitro, A. F., & Marcellin, M. W. (2000). Adaptive reconstruction of phased array MR imagery. Magnetic Resonance in Medicine, 43(5), 682–690. https://doi.org/10.1002/(SICI)1522-2594(200005)43:5<682::AID-MRM10>3.0.CO;2-G

Yacoub, E., Duong, T. Q., Van De Moortele, P. F., Lindquist, M., Adriany, G., Kim, S. G., Uğurbil, K., & Hu, X. (2003). Spin-echo fMRI in humans using high spatial resolutions and high magnetic fields. Magnetic Resonance in Medicine, 49(4), 655–664. https://doi.org/10.1002/mrm.10433

Yousry, T. A., Schmid, U. D., Alkadhi, H., Schmidt, D., Peraud, A., Buettner, A., & Winkler, P. (1997). Localization of the motor hand area to a knob on the precentral gyrus. A new landmark. Brain, 120(1), 141–157. https://doi.org/10.1093/brain/120.1.141

Yu, Y., Huber, L., Yang, J., Fukunaga, M., Chai, Y., Jangraw, D. C., Chen, G., Handwerker, D. A., Molfese, P. J., Ejima, Y., Sadato, N., Wu, J., & Bandettini, P. A. (2022). Layer-specific activation in human primary somatosensory cortex during tactile temporal prediction error processing. NeuroImage, 248(July), 118867. https://doi.org/10.1016/j.neuroimage.2021.118867

Yu, Y., Huber, L., Yang, J., Jangraw, D. C., Handwerker, D. A., Molfese, P. J., Chen, G., Ejima, Y., Wu, J., & Bandettini, P. A. (2019). Layer-specific activation of sensory input and predictive feedback in the human primary somatosensory cortex. Science Advances, 5(5), 1–10. https://doi.org/10.1126/sciadv.aav9053

Yushkevich, P. A., Piven, J., Hazlett, H. C., Smith, R. G., Ho, S., Gee, J. C., & Gerig, G. (2006). User-guided 3D active contour segmentation of anatomical structures: Significantly improved efficiency and reliability. NeuroImage, 31(3), 1116–1128. https://doi.org/10.1016/j.neuroimage.2006.01.015

Zaretskaya, N., Bause, J., Polimeni, J. R., Grassi, P. R., Scheffler, K., & Bartels, A. (2020). Eye-selective fMRI activity in human primary visual cortex: Comparison between 3 T and 9.4 T, and effects across cortical depth. NeuroImage, 220(October 2019). https://doi.org/10.1016/j.neuroimage.2020.117078

