## Supplementary Figures for "Feasibility of 3T layer-dependent fMRI with GE-BOLD using NORDIC and phase regression"

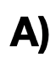

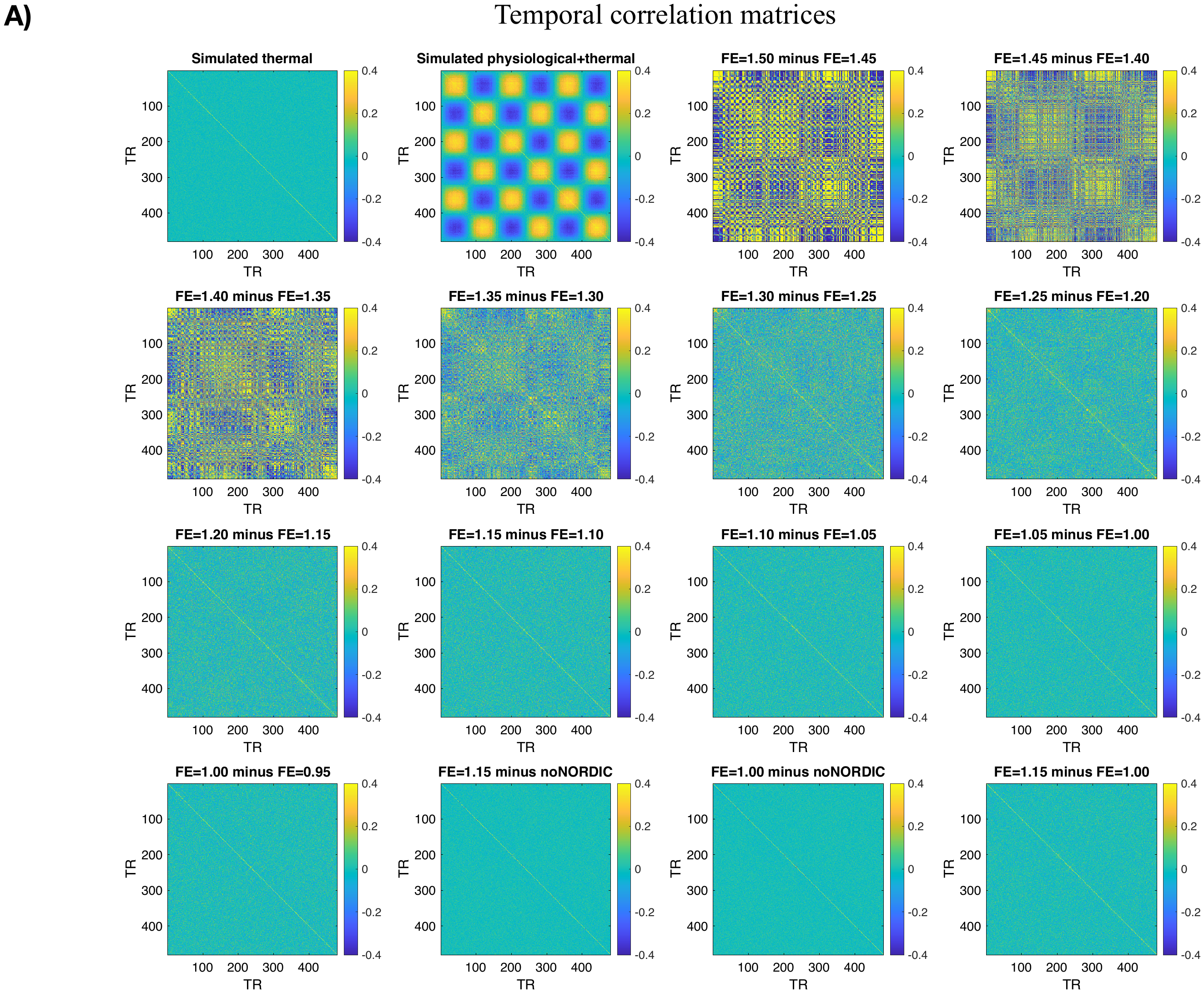

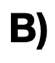

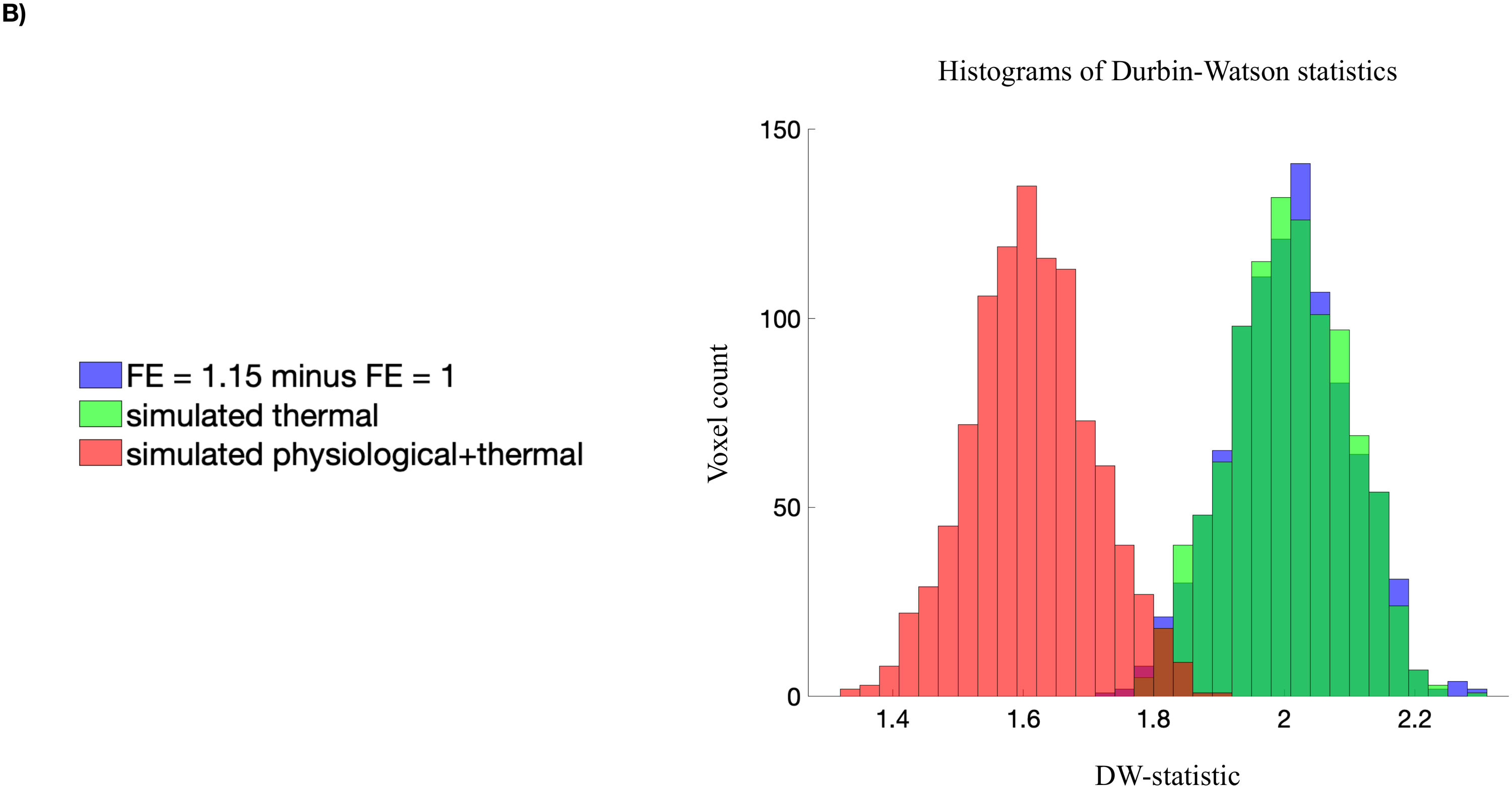

**Figure S1.** Does NORDIC remove thermal noise only? **A)** The first column of the first row (plot 1.1) shows the temporal correlation matrix for 1000 simulated thermal noise timeseries (480 timepoints). Similarly, plot 1.2 shows the temporal correlation matrix for 1000 timeseries of simulated thermal noise plus physiological noise (represented as a sine-wave with 3 periods across 480 timepoints). The ratio of standard deviations between thermal and physiological noise was 2 to 1. Notice in 1.2 that the addition of non-white components leads to structured patterns of high correlations outside the main diagonal. The remaining plots show temporal correlation matrices of differential timeseries between NORDIC versions with varying noise floors adjusted by the ARG.factor_error (FE) option (see Methods section 2.4.1). These timeseries are extracted from a cubic mask containing 1000 voxels placed around the union ROI of an example subject. Plot 1.3, for instance, represents the temporal correlation matrix of the components that are additionally removed when FE is 1.5 compared to when it is 1.45. This difference timeseries clearly contains non-white components. The off-diagonal structure progressively diminishes with lower values of FE and it is largely gone in plot 3.1 indicating that the removed components predominantly constitute thermal noise, even at a FE-value of 1.20. In this study, we used FE = 1.15 to be conservative. Changing FE from 1, as in the original NORDIC version, to a value of 1.15 leads to substantially more noise removal (the across-subject mean tSNR within ROI voxels was 39.47 $\pm$ 5.07 and 22.58 $\pm$ 2.66 for FE=1.15 and FE=1, respectively) and as indicated by plot 4.4, the additionally removed components still appear to be dominated by thermal noise **B)** Histograms over the Durbin-Watson statistics computed from the FE=1.15 minus FE=1 difference timeseries (blue), the simulated thermal noise timeseries (green), and the physiological plus thermal noise timeseries (red) from *A)*. The histogram for simulated physiological plus thermal noise is centered around a value below 2 which is suggestive of successive timepoints being correlated as expected for colored noise. In contrast, the histogram of the difference timeseries is largely identical to that of simulated thermal noise and centered around a DW-value of 2, which is suggestive of no autocorrelation at lag 1. This further supports that thermal noise dominated the components that were additionally removed by changing the FE value, but see Figure S2.

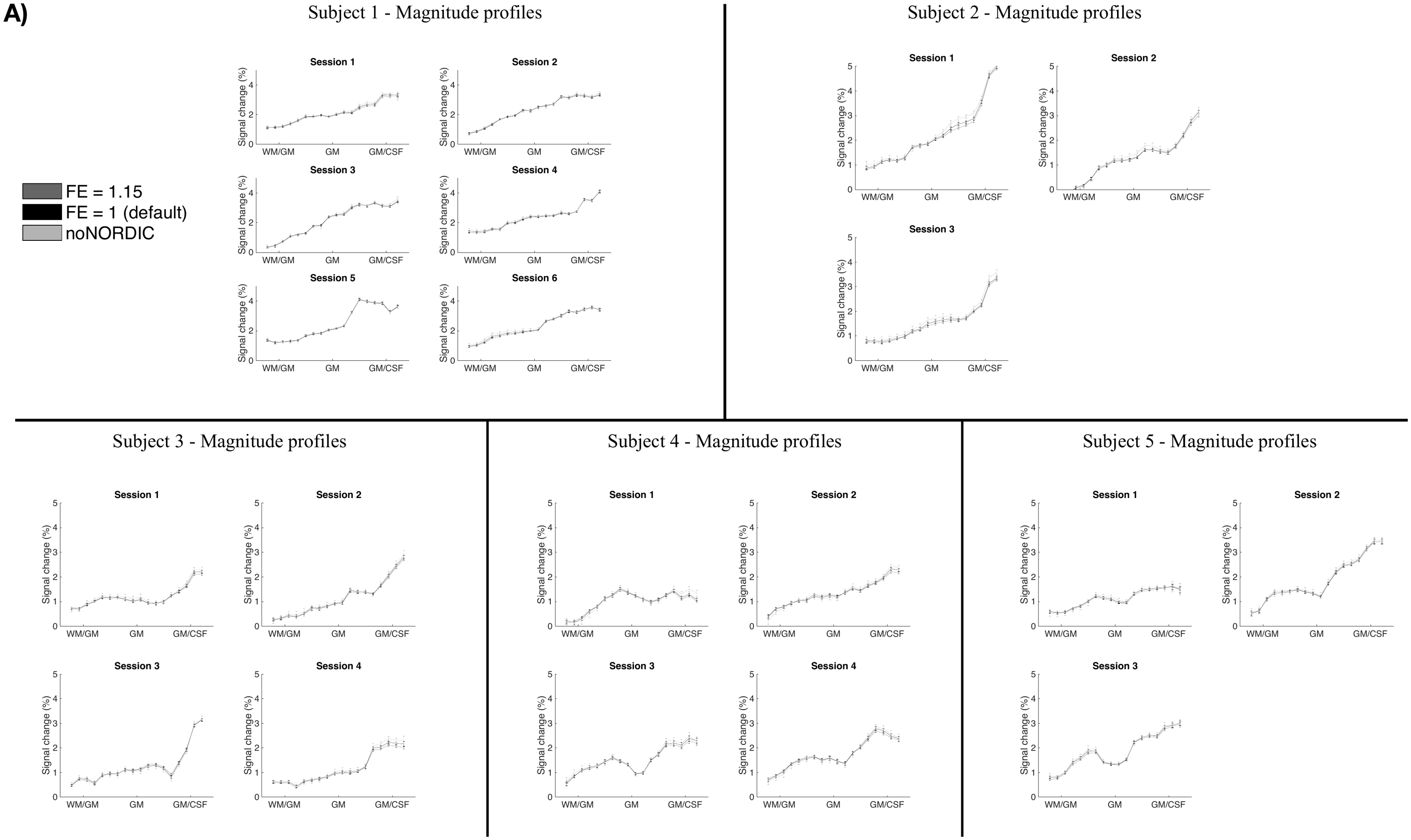

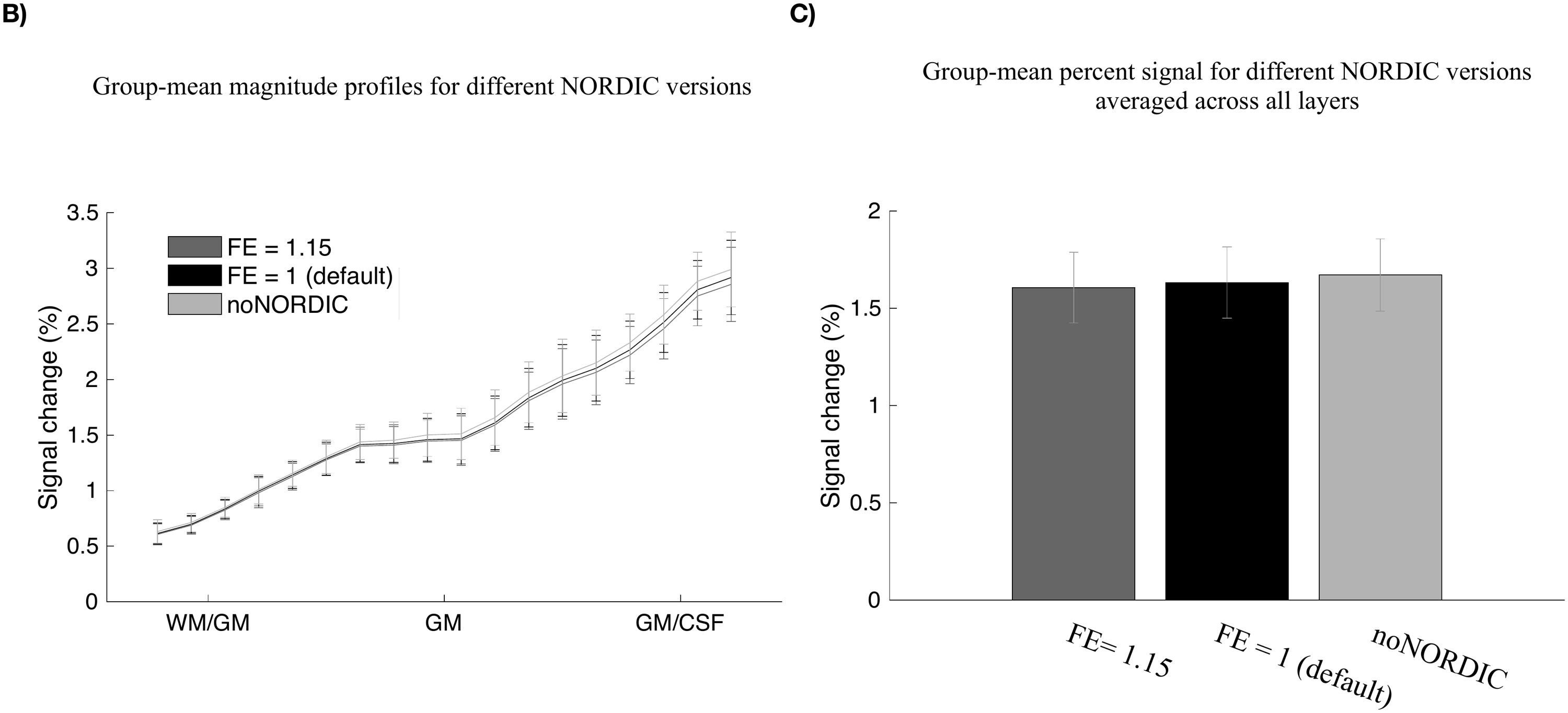

**Figure S2.** Does NORDIC remove signal components? **A)** Magnitude layerprofiles of all sessions and subjects for NORDIC with FE = 1.15, NORDIC with FE = 1, and without NORDIC. Error bars reflect SEM across trials **B)** Across-subject mean profiles for the three NORDIC versions. Error bars reflect SEM across subjects. If NORDIC only removes thermal noise while leaving signal components intact the profiles should be highly similar across NORDIC versions which seems to be the case in both A) and B). **C)** Mean signal change across subjects for each NORDIC version when layers have been pooled by averaging. Paired t-tests (uncorrected) revealed that the signal change of both NORDIC versions was significantly smaller compared to noNORDIC with mean differences of -0.065 percentage points (pp.) $\pm$ 0.039 pp. (p = 0.02) and -0.039 pp. $\pm$ 0.028 pp. (p = 0.037) for FE = 1.15 and FE = 1, respectively. Similarly, the mean signal change for the FE = 1.15 version was significantly smaller compared to the FE = 1 version with a mean difference of -0.026 pp. $\pm$ 0.011 pp. (p = 0.006). This indicates that both NORDIC versions in fact do remove some signal containing components. However, the effect is small and well within the errors associated with the group-mean point estimates and might thus be practically negligible. The magnitude of the effect might differ across setups and the parameter should thus be selected with care in future studies. Notice that all data in this figure is based on beta-estimates for all 30 trials to reduce the impact of noise in the comparison between NORDIC versions.

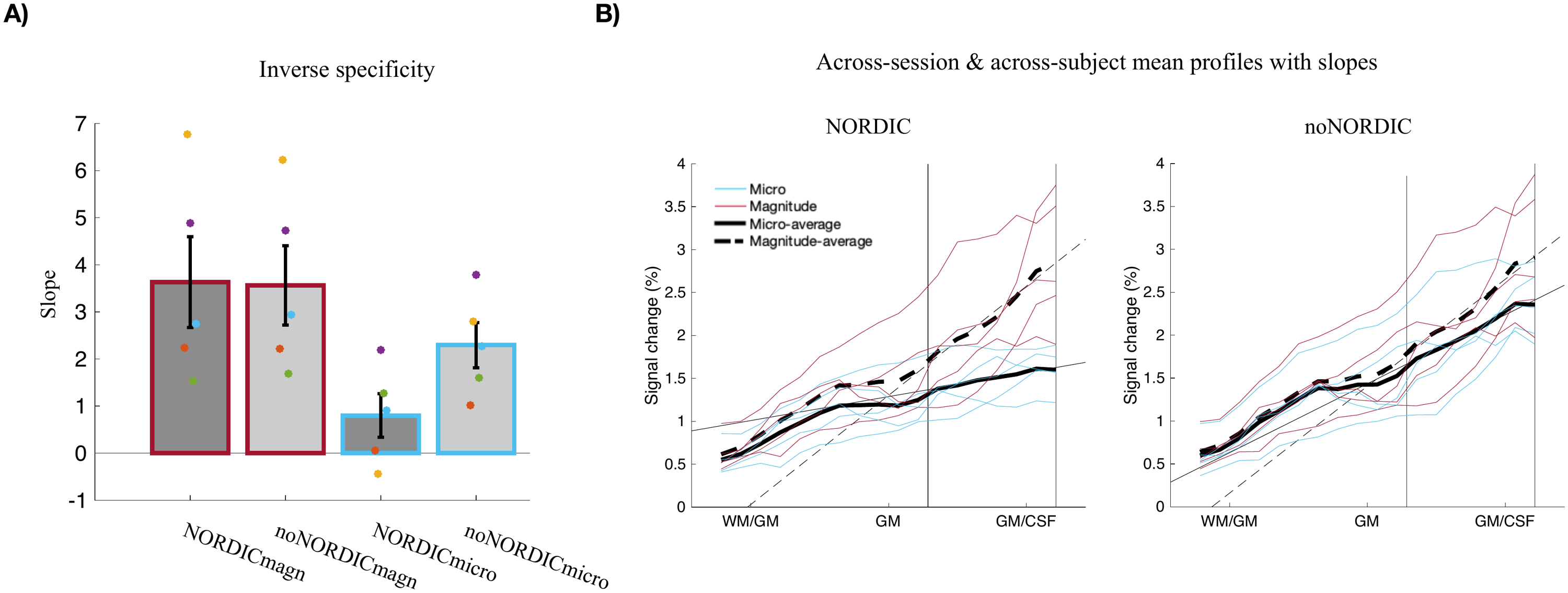

**Figure S3.** Same as Figure 5, except that the slopes were computed between relative depths 0.6-0.925 only (marked by vertical lines). This was done because of the ROI selection criteria that the signal corresponding to relative depths between 0.3-0.4 (deep layers) could not exceed that of relative depths between 0.6-0.8 (superficial layers). The slope between these depth intervals is thus forced to be larger than 0 which might bias the estimated slopes of NORDICmicro profiles towards larger values. Slopes seem to decrease when only the superficial part of the profile is taken into account during the fit as opposed to when the full depth is considered (Figure 5A) suggesting that the ROI selection criteria adopted here might lead to overestimated slopes of the NORDICmicro profiles and thus an underestimated effect of phase regression. Also note, that the slopes of the magnitude profiles seem to increase when only the superficial part of the profile is taken into account suggesting that the characteristic positive gradient of the magnitude profiles from WM towards CSF was not simply an artefact of the ROI selection criteria.
